# The Wound Environment Agent-based Model (WEABM): a digital twin platform for characterization and complex therapeutic discovery for volumetric muscle loss

**DOI:** 10.1101/2024.06.04.595972

**Authors:** Chase Cockrell, Yoram Vodovotz, Ruben Zamora, Gary An

## Abstract

Volumetric Muscle Loss (VML) injuries are characterized by significant loss of muscle mass, usually due to trauma or surgical resection, often with a residual open wound in clinical settings and subsequent loss of limb function due to the replacement of the lost muscle mass with non-functional scar. Being able to regrow functional muscle in VML injuries is a complex control problem that needs to override robust, evolutionarily conserved healing processes aimed at rapidly closing the defect in lieu of restoration of function. We propose that discovering and implementing this complex control can be accomplished by the development of a Medical Digital Twin of VML. Digital Twins (DTs) are the subject of a recent report from the National Academies of Science, Engineering and Medicine (NASEM), which provides guidance as to the definition, capabilities and research challenges associated with the development and implementation of DTs. Specifically, DTs are defined as dynamic computational models that can be personalized to an individual real world “twin” and are connected to that twin via an ongoing data link. DTs can be used to provide control on the real-world twin that is, by the ongoing data connection, adaptive. We have developed an anatomic scale cell-level agent-based model of VML termed the Wound Environment Agent Based Model (WEABM) that can serve as the computational specification for a DT of VML. Simulations of the WEABM provided fundamental insights into the biology of VML, and we used the WEABM in our previously developed pipeline for simulation-based Deep Reinforcement Learning (DRL) to train an artificial intelligence (AI) to implement a robust generalizable control policy aimed at increasing the healing of VML with functional muscle. The insights into VML obtained include: 1) a competition between fibrosis and myogenesis due to spatial constraints on available edges of intact myofibrils to initiate the myoblast differentiation process, 2) the need to biologically “close” the wound from atmospheric/environmental exposure, which represents an ongoing inflammatory stimulus that promotes fibrosis and 3) that selective, multimodal and adaptive local mediator-level control can shift the trajectory of healing away from a highly evolutionarily beneficial imperative to close the wound via fibrosis. Control discovery with the WEABM identified the following design principles: 1) multimodal adaptive tissue-level mediator control to mitigate pro-inflammation as well as the pro-fibrotic aspects of compensatory anti-inflammation, 2) tissue-level mediator manipulation to promote myogenesis, 3) the use of an engineered extracellular matrix (ECM) to functionally close the wound and 4) the administration of an anti-fibrotic agent focused on the collagen-producing function of fibroblasts and myofibroblasts. The WEABM-trained DRL AI integrates these control modalities and provides design specifications for a potential device that can implement the required wound sensing and intervention delivery capabilities needed. The proposed cyber-physical system integrates the control AI with a physical sense-and-actuate device that meets the tenets of DTs put forth in the NASEM report and can serve as an example schema for the future development of Medical DTs.

## 1.0 Introduction to Volumetric Muscle Loss Injury

Volumetric muscle loss (VML) is loss of skeletal muscle following traumatic injury that results in a functional deficit due to the healing of the wound with non-functional fibrotic scar instead of functional muscle [1, 2]. Wound healing of a VML (or more generalized skeletal muscle) injury in mammals is considered to progress in three broad phases: 1) hemostasis, occurring immediately to hours after the injury, 2) inflammation, occurring from hours to days post-injury, and finally, 3) repair, occurring in days to months post-injury [3, 4]. In the hemostatic phase, chemokines, cytokines and other molecular mediators indicating tissue injury are released, which will cause the initial recruitment of leukocytes. Polymorphonuclear neutrophils are the first inflammatory cells to arrive and have been observed as early as one-hour post-injury. [3, 5-7] Endothelial cells release interleukin-8 (IL-8), platelet activation factor (PAF), and a host of other mediators which influence the neutrophils to attach to the endothelial wall. The neutrophils then extravasate through the blood vessel wall and enter the wound, where they eliminate foreign debris and kill bacteria, if present [8]. Subsequently, the neutrophils undergo apoptosis and their pro-inflammatory functionality is replaced by M1-polarized macrophages [3, 9], which also have antimicrobial and phagocytic functions [10].

After this initial pro-inflammatory phase, negative feedback processes lead to macrophage population polarity transitions to the M2 (anti-inflammatory) phenotype, accentuating the suppression of pro-inflammatory processes and causing increased cell proliferation and influx of fibroblasts, extracellular matrix (ECM) synthesis, and capillary budding [11-16]. Over time, depending on the size of the VML injury, fibroblasts differentiate into myofibroblasts, which accentuate wound contraction, the formation of granulation tissue and subsequent conversion to fibrotic scar, marked by the replacement of collagen-3 with collagen-1 [6, 7, 12, 17-19]. The result is a healed, but nonfunctional volume of fibrotic scar tissue existing in place of muscle.

Evolutionary imperatives to enhance the survival of an organism to injury have led to the prioritization of fibrosis and wound closure over the regrowth of functional muscle. A VML injury involves significant loss of tissue and results in a relatively large wound surface area that is exposed to the external environment. To maximize survivability, it is critical to minimize pathogens from the external environment from invading the wound, and thus, the mammalian system quickly fills the missing volume with granulation tissue (i.e., extracellular matrix (ECM), fibrin, fibronectin, etc.) which is ultimately remodeled into stable fibrotic scar tissue [3, 12, 20]. It is this highly evolutionarily conserved, generally beneficial process that must be overridden to facilitate the regrowth of functional muscle in these VML. We propose that this is a complex control problem that requires manipulation of the cellular-molecular biology of skeletal muscle’s response to injury and is suited to the application of a Digital Twin (DT) paradigm.

### 1.1 Medical Digital Twin of VML: Personalization and Precision Medicine

The ultimate goal of biomedical research is to provide the right treatment for the right patient at the right time. We have formalized these principles into the Axioms of “True” Precision Medicine [21]:

- Axiom 1: Patient A is not the same as Patient B (Personalization)
- Axiom 2: Patient A at Time X is not the same as Patient A at Time Y (Precision)
- Axiom 3: The goal of medicine is to treat, prognosis is not enough (Treatment)
- Axiom 4: Precision medicine should find effective therapies for every patient and not only identify groups of patients that respond to a particular regimen (Inclusiveness)

We believe that these Axioms have considerable overlap with the concept of Digital Twins (DTs). The concept of DTs derives from manufacturing and operations fields, including NASA, originating in the 1990-2000s as a means of using computer analogs to real world objects or processes to aid in these products’ lifecycle management, facilitating product monitoring and maintenance. The original formal definition of a DT includes [22]:

- A data structure for the real-world system
- Some process that links data together to form dynamics
- Some link to the real world that feeds back data into the data-propagation/generation process

DTs have found extensive use in industrial applications, both for manufacturing processes and specific engineered systems (e.g., aircraft engines). The key appeal of DTs is that they are “twinned” to a specific real-world object, where the “twinness” between the digital and real-world object is maintained by an ongoing data link between the twins; this allows simulations to be run on the DT to predict the behavior of the real-world twin, anticipating potential failures and proscribing an individualized maintenance plan. Given the rapid and unstructured expansion of the use of the description “Digital Twin,” the United States National Academies of Science, Engineering and Medicine (NASEM) sought to bring some structure and provide guidance by generating an extensive report on this technological approach. NASEM released its report “Foundational Research Gaps and Future Directions for Digital Twins (2023)” after an in-depth evaluation of the potential application of DT to Atmospheric Science, Engineering and Biomedicine [23]. We consider this report (henceforth referred to as the “NASEM Report”) to be an authoritative statement on what constitutes a DT, its unique capabilities and associated research challenges for the promise of the technology to be eventually met.

One of the most common questions regarding DTs is “what is the difference between a DT and a computational model”? The NASEM Report (page 2) addresses this with the following text:

> *“Finding 2-1: A digital twin is more than just simulation and modeling.*

> *Conclusion 2-1: The key elements that comprise a digital twin include (1) modeling and simulation to create a virtual representation of a physical counterpart, and (2) a bidirectional interaction between the virtual and the physical. This bidirectional interaction forms a feedback loop that comprises dynamic data-driven model updating (e.g., sensor fusion, inversion, data assimilation) and optimal decision-making (e.g., control, sensor steering).”*

This definition distinguishes a DT from a personalized predictive computational model, such as an -omics based profile model. which are predicated on a feature set regarding a patient inputted at time of the creation of the model and is not (and often cannot be) updated in an ongoing fashion. We emphasize selected findings from the NASEM Report that directly impact the implementation of a DT to increase the functional recovery from VML. These include:

- Importance of Fit for purpose in defining the requirements of a DT
- Importance of ongoing data-linkage between the real and virtual worlds
- Importance of Establishing Trust in the DT (Validation and Uncertainty Quantification)
- Importance of control as a purpose of the DT

We propose that engineering a solution to the problem of VML will require an adaptive control process that instantiates our Axioms of Precision Medicine using the DT paradigm. Herein we present the computational specification for such a DT, the Wound Environment Agent-Based Model (WEABM). The WEABM is a computational representation of the process of wound healing from a VML injury. The current version of the WEABM is informed by a clinically relevant canine model of VML [24]. In the process of constructing this model, we have integrated knowledge concerning (muscle) wound healing on the volumetric and non-volumetric (e.g., muscle damage/repair from exercise), and have used a calibrated model to suggest broadly defined therapeutic strategies to increase functionality/volume on functional muscle in wounds healing from VML injuries.

### 1.2 Fit-for-purpose and overall description of WEABM

We applied our previously developed schema for determining the design criteria for a VML DT and its computational specification the WEABM [25]. The key features identified are:

- Fit for Purpose: Goal is to identify means of increasing muscle regrowth in VML by manipulation of cellular behavior via molecular-level interventions.
- Primary Scale of Representation: Cellular behavior, aggregated into system-level phenotype (e.g., the VML wound).
- Data Linkages: Hypothesized, based on requirements for effective control.

These results led to the decision to construct the WEABM as an Agent-based Model (ABM). Given the clinically relevant anatomic scale of the entire system, the WEABM was implemented in C++ to facilitate execution on high-performance computing platforms. Agent-based modeling is an object-oriented, discrete-event, rule-based, spatially-explicit, stochastic modeling method in which agents interact with each other and with their environment according to a pre-defined set of rules. In this work, the rules are informed by both literature and experimental data. Literature review has been used to establish the existence of a rule, e.g., interleukin-13 (IL13) stimulates fibroblasts to produce Transforming Growth Factor β1 (TGFβ1) [18], and the experimental data is then used to quantify the contribution/effect of IL13 on fibroblast production of TGFβ1 in the model. We note that parameter values of this type are inherently unitless and not directly translatable to the real world (see ‘Model Calibration’ in Methods). In this sense, an ABM is a computational/dynamic instantiation of mechanistic knowledge that has been established through rigorous and reproducible biological experiments. Because ABM’s synthesize knowledge outside the domain expertise of a single researcher, behaviors generated by the aggregate behavior of the agents in the model can often be non-trivial or unintuitive, rendering ABM an excellent technique to investigate biomedical systems. The utility of ABM’s spans from general purpose anatomic/cell-for-cell representations of organ systems capable of reproducing multiple independent phenomena [26, 27] to platforms for drug development,[28, 29] and are frequently used to model non-linear dynamical systems such as the human immune system [30-33]. For this initial implementation, the WEABM is calibrated to a canine model of VML. This model was chosen because it: 1) can produce a large enough wound such that it is contextually clinically relevant (12 cm x 7 cm x 4 cm) in the thigh, 2) allows for sequential tissue sampling so that tissue mediator time series can be obtained to capture the dynamics of the VML response, and 3) allows for experiments to be potentially performed to evaluate potential control strategies aimed at increasing the amount of functional muscle in the healing of the VML injury.

In the WEABM, the individual agents are cells (see Figure 1 for a wire-frame schematic of the cellular-molecular interactions implemented and see Appendix A for “Facts and References for the WEABM”) and the environment is discretized and represented as a three-dimensional grid of cubic voxels measuring 100 μm per side, as this is comparable to the diameter of a type-1 skeletal muscle fiber in the quadricep of a female German Shepherd [34, 35]. Each tissue voxel has three primary data elements associated with it: 1) volume of various tissues: each voxel can be filled with some combination of healthy muscle (which requires adjacent healthy muscle), collagen-3 (broadly representing granulation tissue), and collagen-1 (broadly representing fibrotic scar); [7, 19, 36-39] 2) a Boolean variable indication vascularization, which requires vascularization of a contiguous voxel; and 3) a Boolean variable indicating innervation, which required innervation of a contiguous voxel. Muscle fiber orientation is represented as a restriction on the movement of satellite cells such that, in the undamaged tissue, muscle fibers are oriented along the y-axis of the simulation and satellite cells are myoblasts are required to migrate along a single myofiber. In damaged regions, satellite cells and myoblasts are allowed free movement [5, 6, 40].

**Figure 1:**
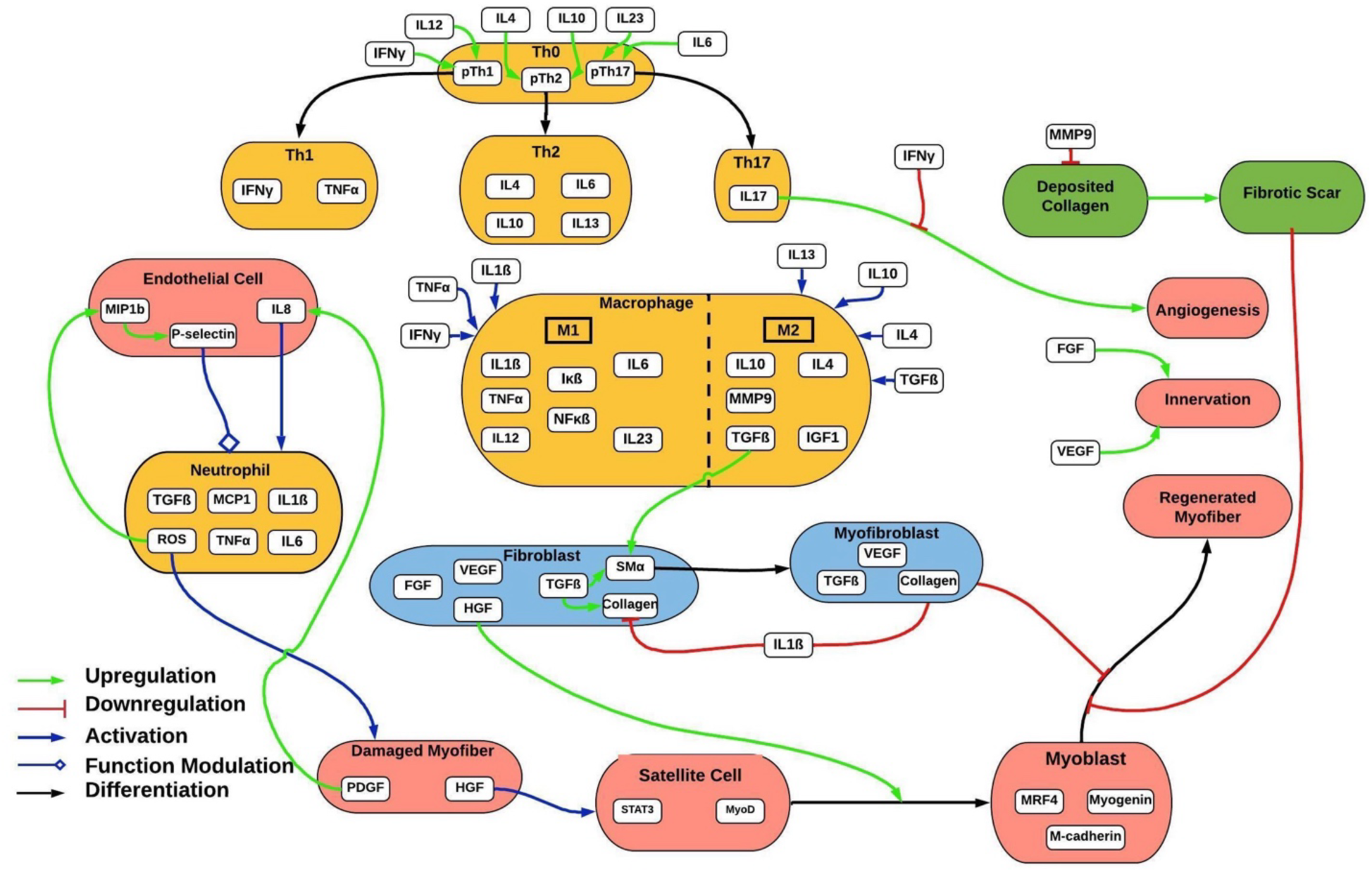
Schematic of the WEABM Immune cells are orange boxes, muscle regeneration cells and neurovascular components are depicted in pink, scar-promoting cells in blue and extracellular matrix/collagen in green. Green arrows denote promoting interactions, red denote downregulation blue represents upregulation and black represents differentiation.

An execution of the WEABM proceeds by first instantiating the wound, and then incrementing the simulation in time. Each time step of the simulation represents 15 minutes. At the start of each step, the different cell types are randomly shuffled, and each cell then performs its actions sequentially. After the cells perform their actions, the diffusion and degradation of mediators and cytokines are simulated. Cells that have died (from apoptosis or necrosis) are then removed from the simulation and any new cells generated from proliferation or division are added to the simulation world.

The WEABM can be run in two primary modes: 1) the full anatomic scale that corresponds to the 12 cm long x 7 cm wide x 4 cm deep experimental wound in the reference canine model and can represent up to 400 million cells or 2) a biopsy scale that represents a smaller volume (approximately 6 mm long x 6 mm wide x 6 mm deep) corresponding to the size of tissue biopsies in the reference canine model [24]. In the anatomic scale configuration initial conditions are generated through processing data obtained from a CT scan of the canine model immediately after the application of the surgical wound; this allows for personalization of the WEABM to a specific individual consistent with the digital twin paradigm. We first transform the surface to remove the curvature of the dog’s leg, as described in [41], which essentially projects the wound into a Euclidean space with the surface of the wound at the top of a rectangular prism and the depth of the wound going through the z-axis (See Figure 2).

**Figure 2:**
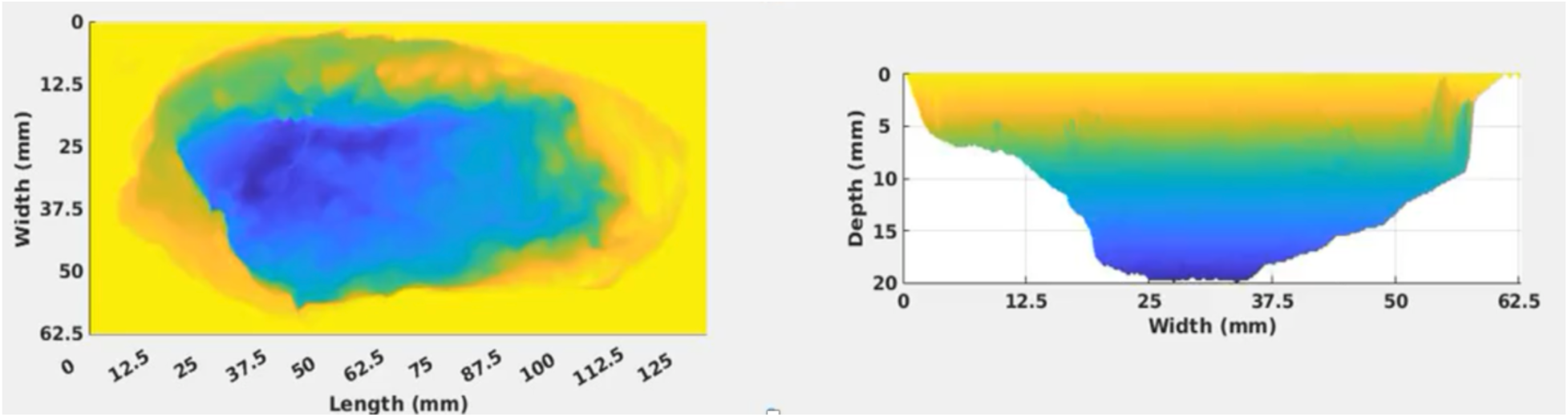
Heatmap of XYZ Axis representation of the WEABM with initial conditions mapped from the first post-surgical Computer Tomography (CT) scan of a canine wound. Color scale is related to the depth of the wound (blue deep and yellow superficial). Tissue is shaded based on distance from the ideal wound surface, shaded from yellow (superficial) to blue (deep). The panel on the left shows a top-down view of the wound (looking down the z-axis at the x-y plane) and the panel on the right shows a side view (looking along the x-axis at the y-z plane).

We consider all tissue that is exposed to the atmosphere (and all tissue below that along the z-axis) to be healthy muscle. Given the fact that one of the hallmarks of acute VML is that the sizes of the defects preclude closure of the wound with biological tissue, we recognize that the surface of the wound is exposed to the atmosphere/environment and either that atmospheric exposure and/or any dressing/wound device applied to that surface will generate an ongoing inflammatory stimulus that would initiate and propagate the injury response process. A rendering of a personalized initialization of the anatomic scale WEABM can be seen in Figure 3.

**Figure 3:**
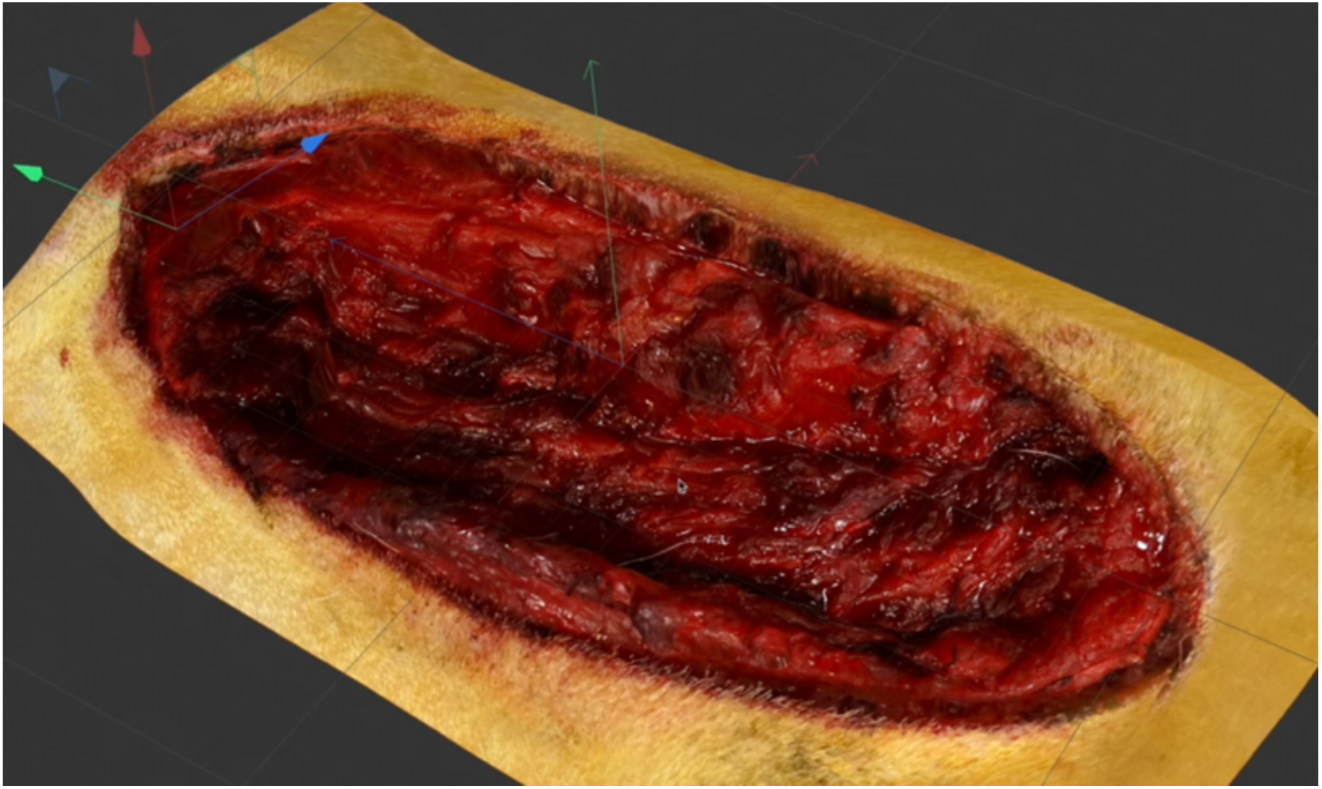
Rendering of a personalized initialization of the WEABM. This image adds realistic tissue textures to the underlying voxel structure of the WEABM. Rendering produced by 4dThieves™.

In the biopsy configuration, the WEABM simulates a volume of tissue which is comparable to the experimental biopsies, specifically, a 6-mm (60 voxels on each side) cubic section of tissue represented with a rectangular prism of tissue. The WEABM in the biopsy configuration can grow to a larger depth than 6mm, however only the top 6mm of tissue are used to compare protein concentrations with the experiment and the very top of the Z-axis represents the wound-atmosphere interface noted above. Further, there is an additional abstraction here regarding the location of satellite stem cell pools. In the anatomic configuration, each muscle fiber has at least one satellite stem cell pool randomly located in the healthy (uninjured) tissue at the boundary of the simulation; in the biopsy configuration, the only healthy tissue is located at the base of the tissue volume, and thus, we allow myoblasts to initiate new fibers at one edge of the simulation-space.

There has been extensive prior work using multiscale mechanism-based modeling to study skeletal muscle injury and recovery, both using finite element methods [42, 43], and ABMs [44-46], including work using an ABM specifically examining and proposed means of improving muscle regrowth in VML [47]. The WEABM has many similarities to the work of Westman [47], particularly in terms of the inclusion of the primary cell types involved and the molecular signaling pathways present in VML. Primary features that distinguish the WEABM from prior work include: 1) the fact that the WEABM is a 3-dimensional model, 2) the WEABM is capable of representing clinically relevant, anatomic scale injuries and 3) the representation of the WEABM incorporates a critical pathophysiological feature seen in clinical VML, namely that the wound is not covered with biological tissue and therefore the wound-environment interface represents an ongoing injury/inflammatory source in VML.

To demonstrate the potential utility of the WEABM as the basis of a VML DT, we performed a series of simulation experiments with the WEABM with the goal of governing a complex adaptive control process that would increase the recovery of functional muscle in VML. The simulation experiments include:

- Initial Evaluation of Behavior Space of the WEABM to provide fundamental insights into the healing biology of VML
- Initial Calibration to Tissue Protein/Mediator Data to fulfill the validation requirement noted in the NASEM Report.
- Perform control discovery using simulation-based Deep Reinforcement Learning (DRL)

- Simulated Control Modalities:

- Multimodal mediator administration
- Application of Extracellular Matrix (ECM)
- Combination multimodal therapy including a hypothetical anti-fibrosis agent

The results of these simulation experiments are provided below. The code for the WEABM and the DRL environment can be found at https://github.com/An-Cockrell/WEABM_DRL/tree/main.

## 2.0 Results

### 2.1 Pre-Calibration Insight: The essential role of the competition between fibrosis and myogenesis

Initial simulation experiments explored model behavior space arising from the inherent model structure of the WEABM. These simulations provided fundamental insights into the dynamics that governed healing of VML, and therefore served to frame subsequent investigation of potential control strategies aimed at healing these wounds with potentially functional muscle. The first of these insights was the essential role of the competition between fibrosis and myogenesis. Figure 4 shows a set of early pre-calibration simulations that examine the behavioral range of the WEABM.

**Figure 4.**
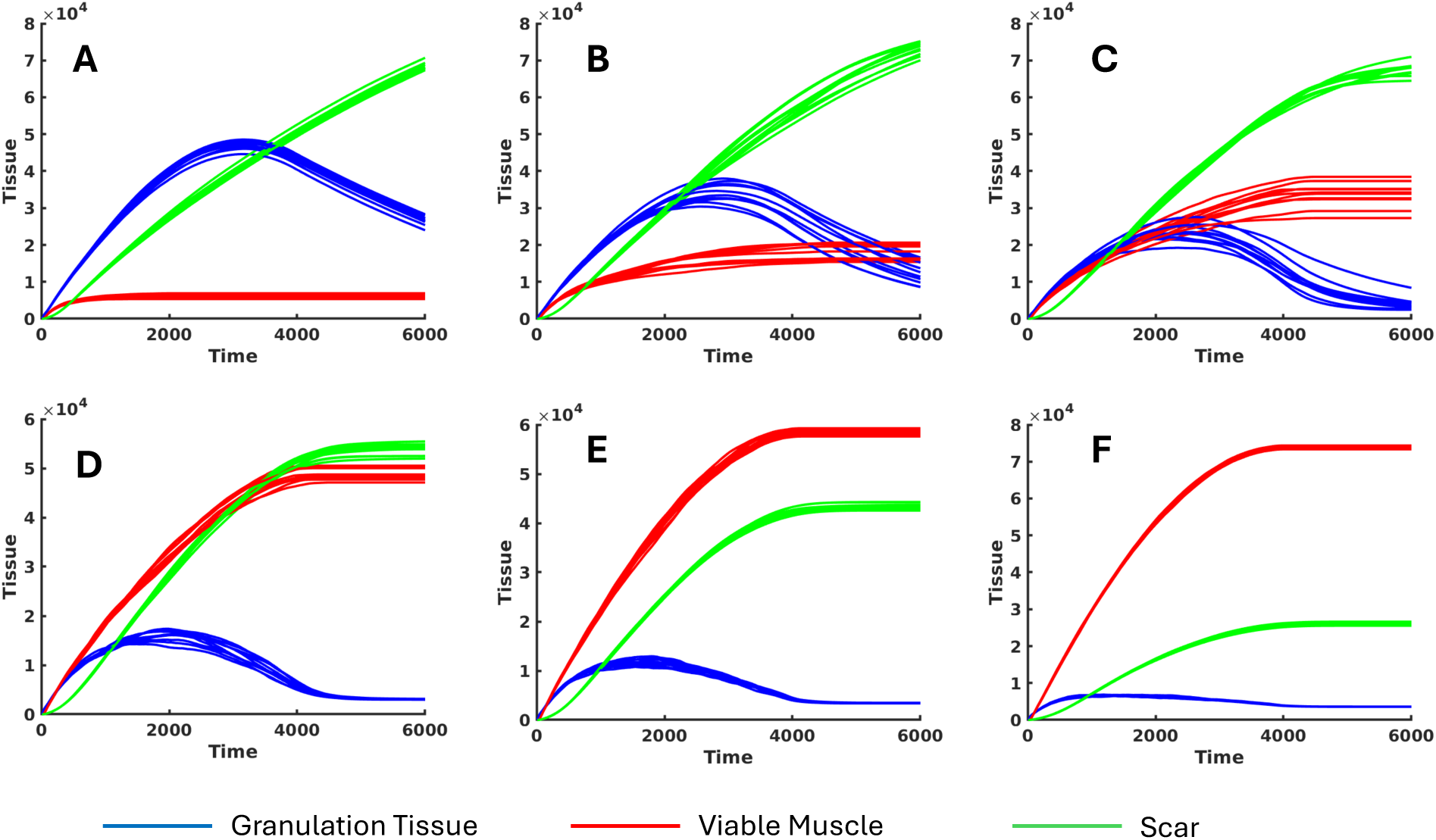
Parameter space exploration demonstrating competition between fibrosis and myogenesis. With adjustment of parameters governing the rate of fibrosis functions and myogenesis functions, a progression of tissue composition during healing can be seen in the progression of Panel A through F, where Panel A represents the “native” configuration where fibrosis predominates, Panels B-E depict a shift between fibrosis and myogenesis, towards Panel F, where effective muscle regrowth is generated. Though uncalibrated, these dynamics show a potential path towards the desirable outcome of increased functional healing.

Specifically, we focused on the processes associated with fibrosis and myogenesis. The model structure demonstrated that there is a fundamental tradeoff between these two processes, a competition for binding sites on the edge of the damaged health muscle where myoblasts would fuse to differentiate into functional myofibrils. We term this competition “the race between fibrosis and myogenesis,” and this dynamic informs our subsequent calibration simulations and the process of control discovery.

### 2.2 Calibration and Validation

This first step in calibration is determining if the overall healing dynamics of the canine VML can be reproduced. These results can be seen in Figure 5, which compares the wound closure pattern between experimental data and WEABM simulations.

**Figure 5.**
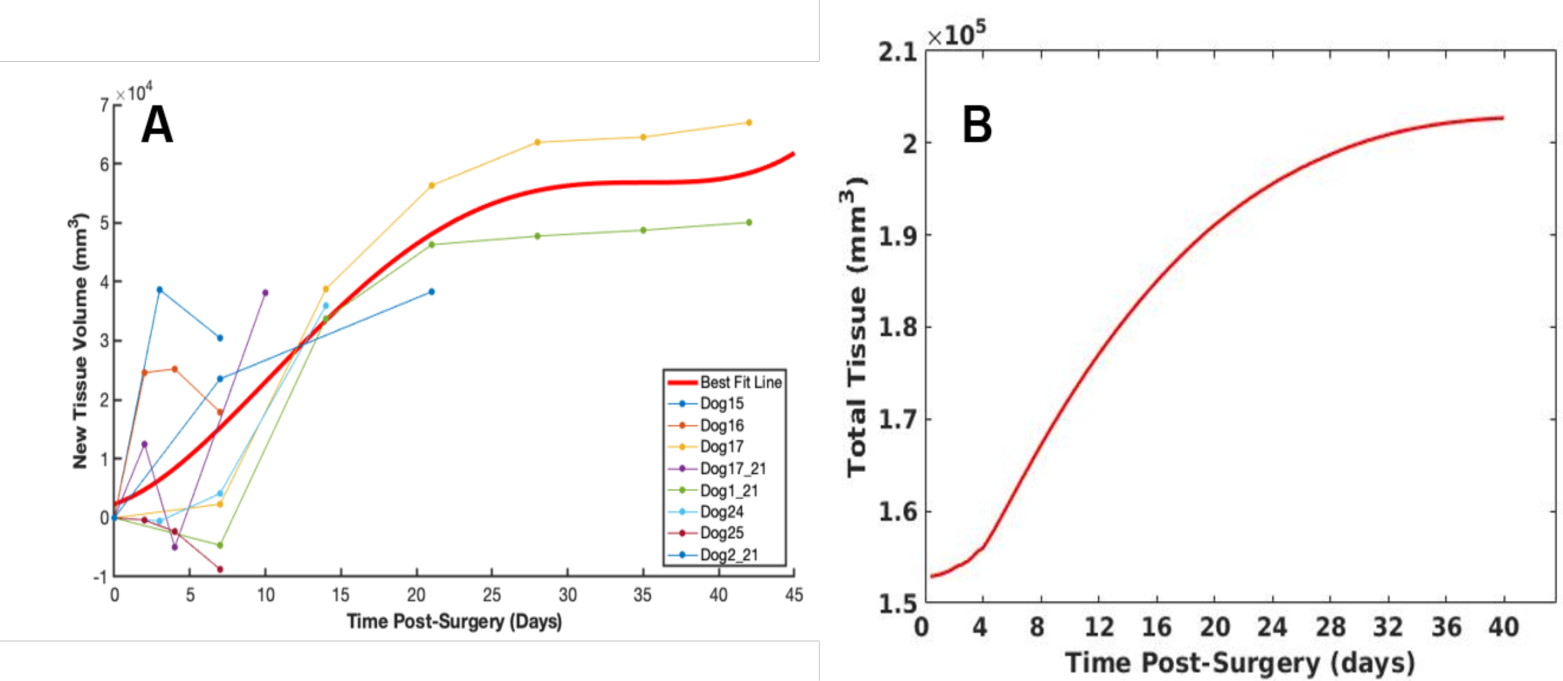
Comparison of tissue growth. Panel A: Total new tissue growth in each dog calculated from CT data, as well as the best-fit line in red. We note that there are many sources of experimental error when obtaining this experimental measurement, including dog motion and tissue swelling, and as such, we posit that the best-fit line is representative of true tissue growth. Panel B: Total tissue in the volume simulated as the wound evolves over 40 days.

The visual depiction of this process is seen in Figure 6, which shows successive renderings of the VML wound from a specific experimental animal as it heals.

**Figure 6.**
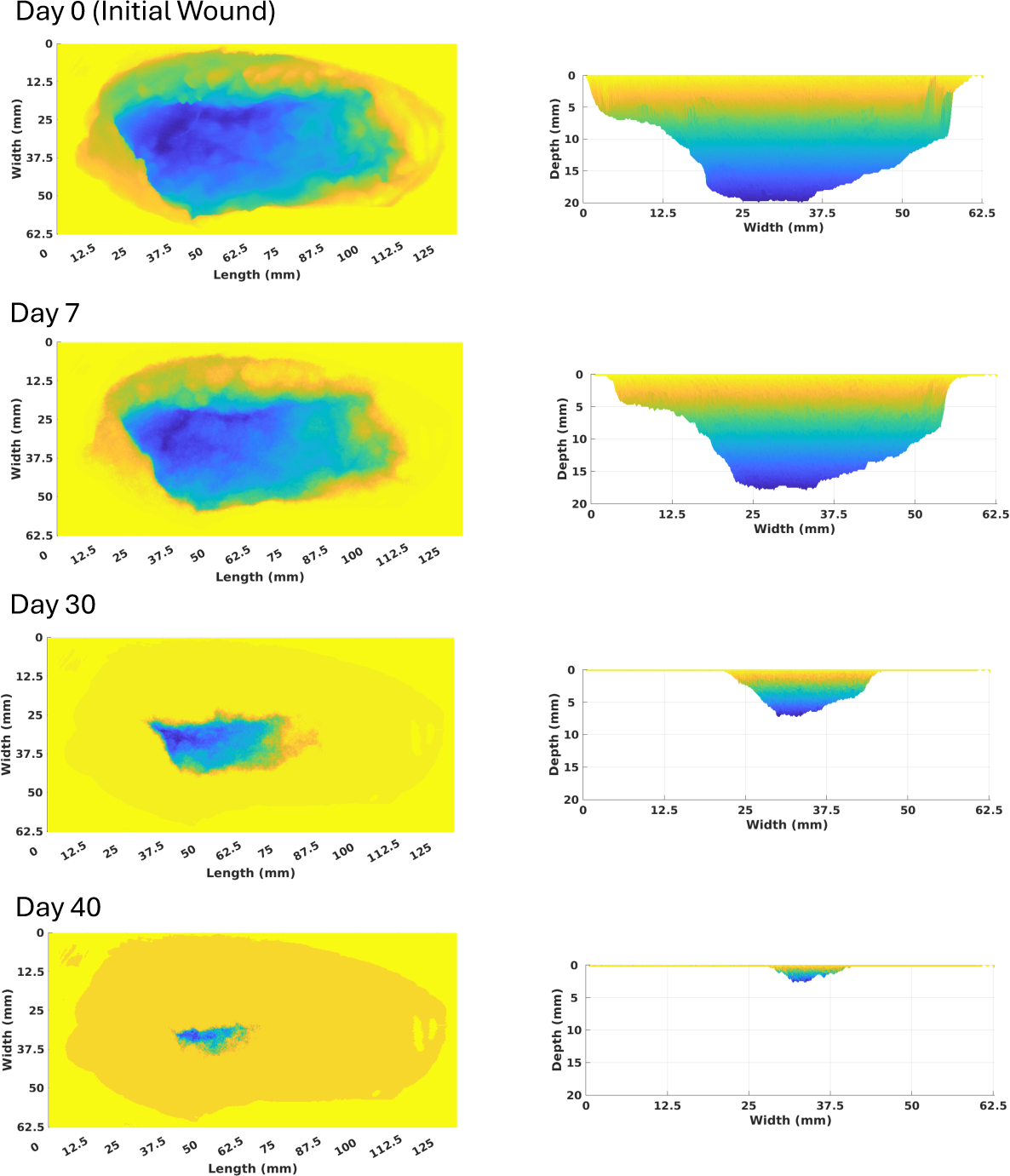
Selected images of the WEABM as it heals from a VML over 40 days simulated time (Days, 0, 7, 30 and 40). Each panel displays a representation of the wound surface only (not the bulk tissue) for rendering reasons. Tissue is shaded based on distance from the ideal wound surface, shaded from yellow (superficial) to blue (deep). The panel on the left shows a top-down view of the wound (looking down the z-axis at the x-y plane) and the panel on the right shows a side view (looking along the x-axis at the y-z plane). Note that since this represents baseline biology the majority of the wound is filled with scar.

Having demonstrated face validity by reproducing the overall healing dynamics of VML injury the next step in calibration is the fitting of the WEABM to experimental time series data of mediator levels in the wound. Our novel process to complex model calibration and validation is described in the Methods, and involves the identification of all model structure and parameter configurations of the WEABM that are sufficient to reproduce the range of the experimental data using a mathematical object called the Model Rule Matrix (MRM) [48, 49] and a machine learning (ML) search pipeline that includes genetic algorithms (GA) and active learning (AL) that we term Nested Active Learning [50]. The result of this process is a set of model structure and parameter configurations that we term *bioplausible* insomuch they cannot be falsified by any available data. As previously noted, the calibration data are drawn from Ref [24]. The results of this process are seen in Figure 7.

**Figure 7:**
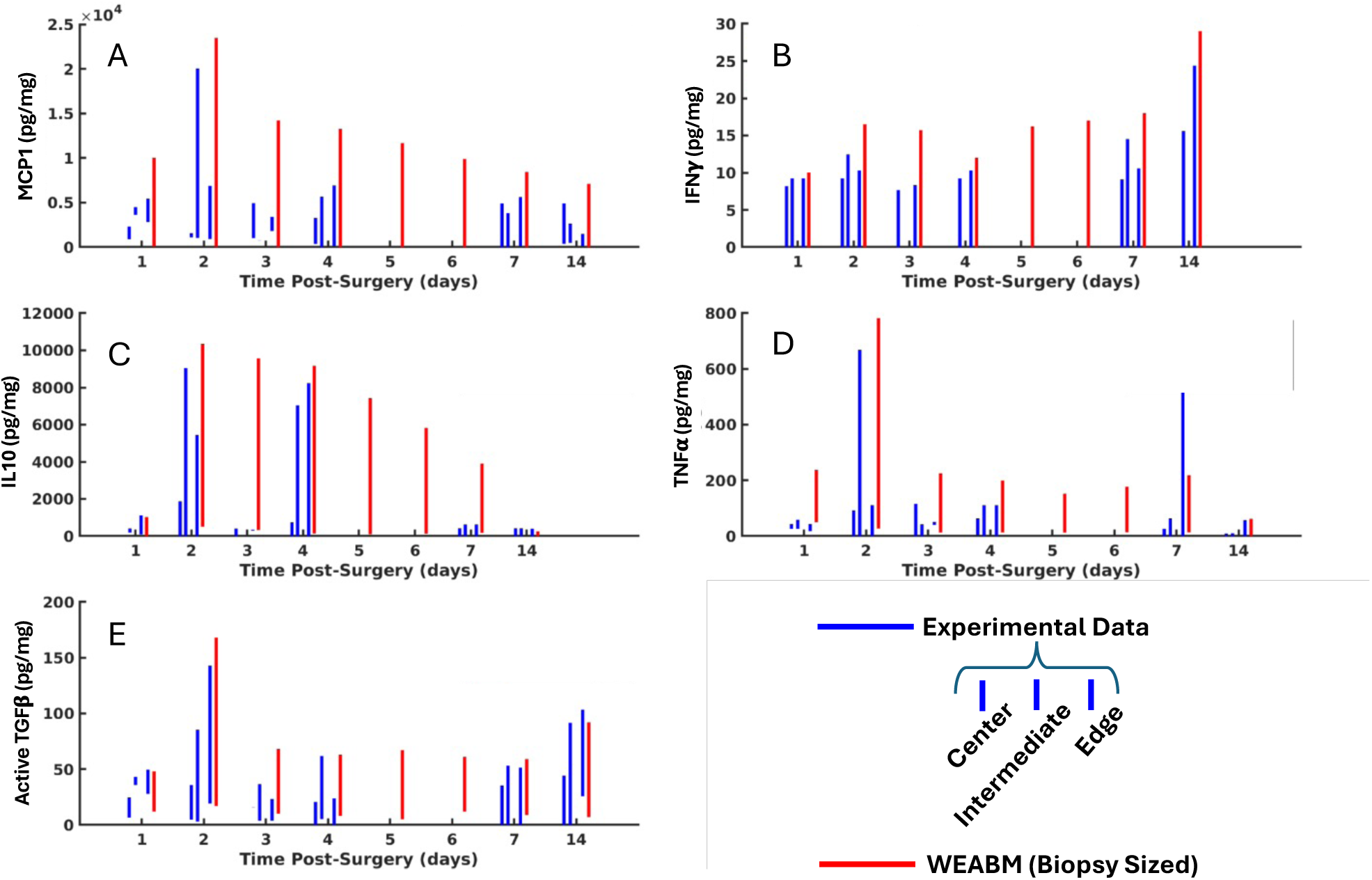
Comparison of WEABM bioplausible output to experimental data from different zones of wound for representative mediators MCP-1 (Panel A), IFNγ (Panel B), IL-10 (Panel C), TNFα (Panel D) and active TGFβ (Panel E). Blue bars are experimental data based on location within wound; the different wound zones are shown for demonstration purposes, with the center zone as the left-most blue bar, the intermediate zone as the center blue bar, and the edge as the right-most blue bar for each day, as the data across these regions are aggregated for determination of bioplausible trajectories of the WEABM (since all regions demonstrate values reachable/bioplausible in the real-world system). Red bars are simulation output of the WEABM at corresponding time points; note that WEABM simulated data projects what values would have been at Day 5 and 6.

We further define bioplausibility as model behavior being within the bounds of the experimental data while maintaining as much behavioral heterogeneity as possible; this is intended to mimic the ability of a particular model configuration to reproduce the breadth of behavior seen in a population, thereby providing generalizability as a property of the model representation. We also note that the Nested Active Learning fits to all selected variables (e.g., multidimensional fitting) at the same time (in this case all the measure mediators in the WEABM, of which 5 are shown in Figure 7). Parameter determination currently equally weighs every mediator for which experimental data exist, therefore there is differential quality of calibration in the 5 mediators shown here. For the most part, the experimental data are contained within the range of ensemble-simulation generated data, with the exception of the spike in TNFα at day 7. Deeper investigation into these measurements identified that they derived from a single animal that was experiencing systemic inflammation due to causes unrelated to the experimental VML and therefore likely had increased systemic levels of TNFα. This finding, rather than showing a limitation of calibration through Nested Active Learning, demonstrates the ability of the process to determine the bounded expressiveness of the underlying model structures as reflected by the set of bioplausible MRMs and can even suggest alternative explanations for outlier measurements. This set of bioplausible MRMs now is the superset of WEABM configurations that are operated on in subsequent simulation experiments.

The next step in validation is the ability of the WEABM to provide forecasting of individual trajectories. The range and heterogeneity in the data set is considerable, as would be expected from a population of non-genetically identical animals (or a clinical population) undergoing a complex surgical procedure. It is impossible, therefore, to attempt to pre-specify where in that entire possible trajectory space an individual animal’s data will reside. However, the DT paradigm, with its ongoing data link between the real world and the digital object, inherently accommodates the concept of a *rolling forecasting cone*, wherein, given a particular point in system state space, a range of possible trajectories can be generated using any ensemble of MRM configurations able to encompass that particular set of multidimensional data points. This cone is then updated with additional data feeds from the real world and propagates forward in time. This is the same concept used in forecasting hurricane paths. The capability of the WEABM is demonstrated in Figure 8, which shows a refined forecasting cone generated by a defined ensemble of MRM configurations for an individual animal as a subset of the total possible trajectories.

**Figure 8.**
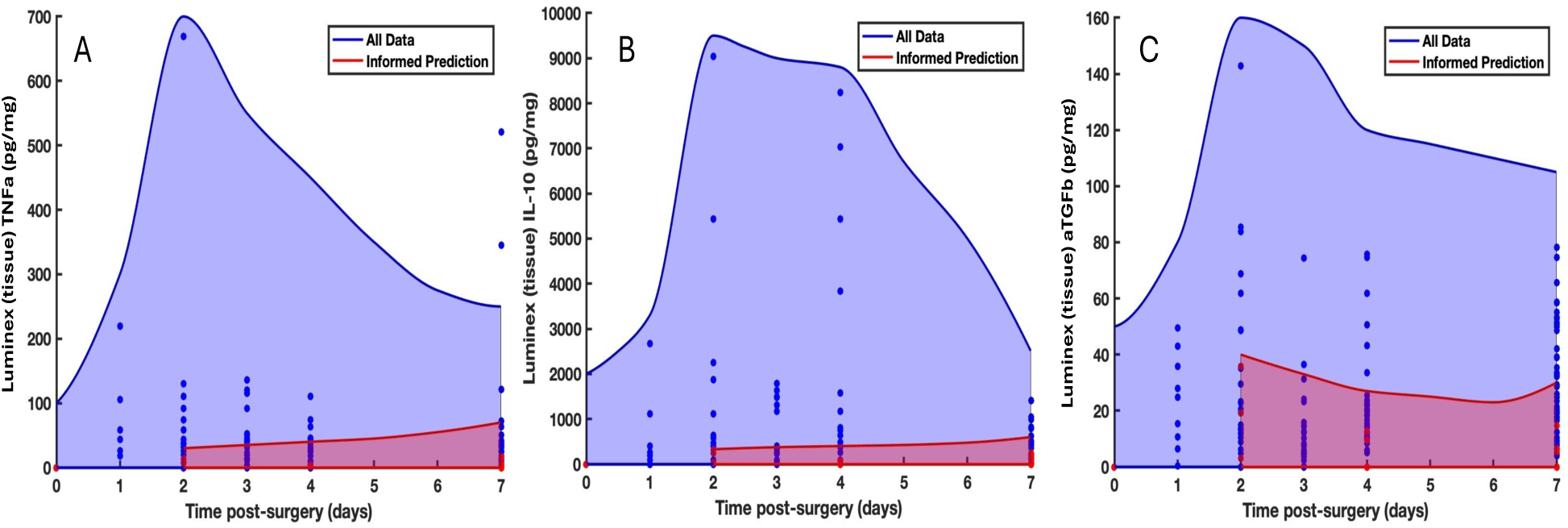
Demonstration of prediction cone for an individual animal. Trajectory spaces are shaded areas, actual experimental measurements are shown as dots. Blue Dots: Experimental data points from all dogs (N = 16). Blue Shaded Area: Total possible trajectory space generated by WEABM (encompasses all Blue Dots). Red Dots: Data points from an individual animal. Red Shaded Area: Forecasting cone for the individual animal, with re-fitting based on updated data. Panel A = TNFα, Panel B = IL-10, Panel C = active TGFβ. The refinement of the forecasting cone over time is analogous to the rolling prediction seen in hurricane prediction, and consistent with the DT paradigm in terms of an updating data feed.

The results of the calibration and validation simulations provided us a baseline level of trust that the WEABM plausibly reproduced critical features of the real-world VML model in terms of overall wound healing dynamics, reproduction of the heterogeneity and trajectories of mediator levels and the ability to individualize/personalize an ensemble of model configurations to a specific individual. As noted in the defined fit-for-purpose of the WEABM-based VML DT, we now turn to exploration of potential control modalities that would be able to increase the amount of functional muscle in the VML wound.

### 2.3 Examining Control Modalities

#### 2.3.1 Insights into control components

Simulation experiments with the WEABM during the model exploration and calibration phases provided insights into the key dynamic features that would be needed to be targeted with putative control strategies. These insights included:

1. Confirmation of importance of competition between fibrosis and myogenesis (see Figure 4). Once collagen deposition occurs, caps the ends of the myofibrils, no myoblast fusion can occur, therefore no muscle regeneration. We tested this hypothesis by arbitrarily enhancing myoblast function to a 4-fold increase in proliferation. The results of this simulation can be seen in Figure 9.
2. Importance of minimizing the duration of the pro-inflammatory/M1 phase, but also recognizing that the anti-inflammatory/M2 phase needed to suppress pro-inflammation was itself detrimental through its pro-growth, pro-fibrosis functions. Specifically, the M2 phenotype conversion of fibroblasts to myofibroblasts and subsequent deposition of early Type 3 collagen led to the evolutionarily favored circumstance where fibrosis would invariably win the race between fibrosis and muscle regrowth.
3. The resulting control paradox: The M2 phenotype was needed to curtail the pro-inflammatory phase but counter-productive to the therapeutic goal of muscle regrowth.
4. The open nature of the VML wound provides an ongoing pro-inflammatory stimulus to the surface of the wound through atmospheric/environmental exposure, essentially providing an ongoing perturbation that “resets” the injury clock, enhancing the amplitude and duration of the pro-inflammatory phase of wound healing (M1) and a subsequent augmentation of the pro-growth/fibrosis processed generated by the compensatory M2 response. The robustness of the fibrosis response arises from the evolutionary imperative to biologically close the wound.

**Figure 9:**
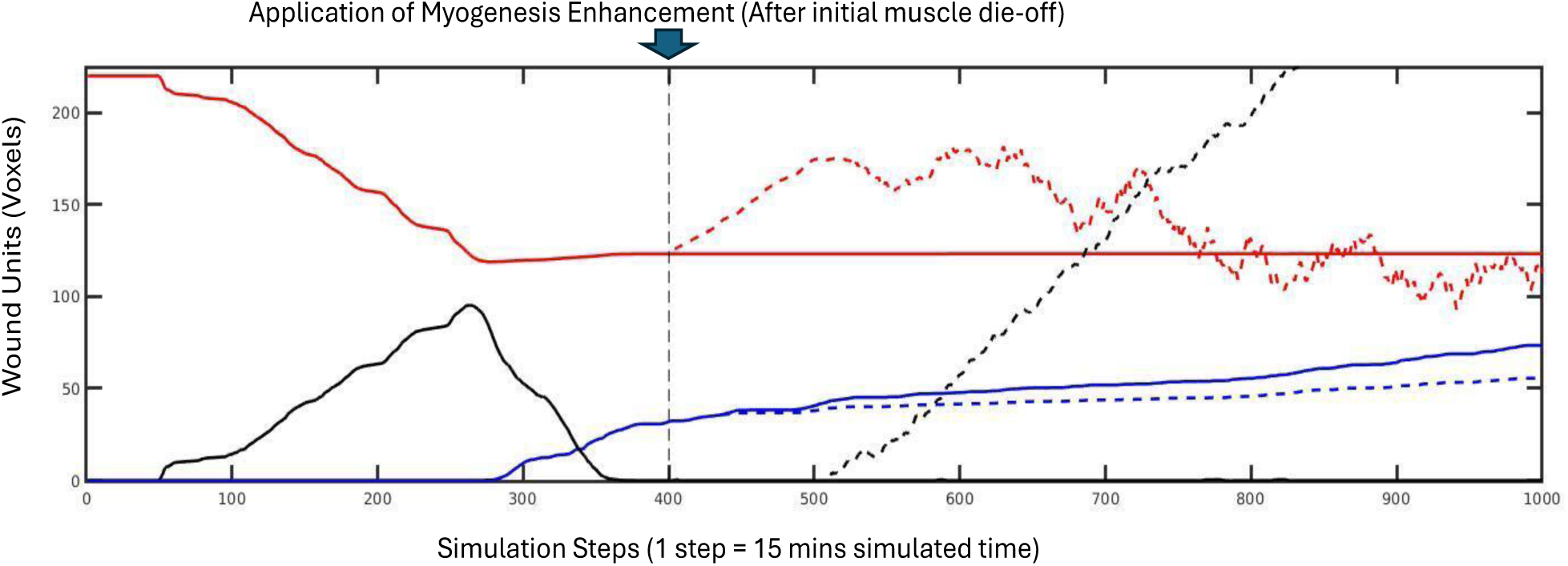
Simulation of increased myogenesis alone. Red lines = healthy muscle, Black Lines = dead muscle, Blue Lines = scar, Solid = Base case, Dotted = 4x increased satellite cell proliferation. Enhanced earlier myoblast function (vertical dashed line is initiation of intervention) implemented as a 4-fold increase in satellite cell proliferation did not increase muscle regrowth and instead increased necrotic “muscle.” Interpretation of these results is that increasing numbers of satellite cells with a fixed number of exposed myofibril ends led to increased numbers of un-attached myoblasts that subsequently died.

All these features point to a complex control problem that can generally be categorized into three main phases:

- Early (up to ∼ 3 days): emphasizing the efficacy and subsequent plateau of the initial pro-inflammatory/M1 response
- Transition to Growth (∼7-14 days): key interval where the initial pro-inflammatory response is attenuated (if possible) and pro-fibrosis functions are initiated as a result.
- Late (after ∼ 14-21 days): in this phase the initial collagen deposition has occurred, and subsequent remodeling would require degradation of existing collagen to re-expose myofibrils for fusion to new myoblasts or rests of differentiated muscle tissue.

We identify the key transition point as being in the 7-10-day period, and therefore posit that this period should be targeted by mediator manipulation to change the relative strengths and efficacy of fibrosis versus myogenesis pathways to shift the conditions of the race between these two modules. Given the known heterogeneity seen in the tissue and the dynamic nature of the processes we propose using the WEABM in a DT context to provide an adaptive sense/actuate control approach to the processes in this phase. More specifically, since this transition phase represents a balance between pro-inflammation (nominally M1), appropriate inflammation suppression via anti-inflammatory mediators (nominally M2), and pro-growth factors associated with the anti-inflammatory processes, we pose that the ability to categorize the wound requires: 1) the ability to sense the pro-inflammatory state of the wound and 2) the ability to sense the canonical anti-inflammatory/pro-growth mediator milieu. These sensing targets will be successively refined in the results of the deep reinforcement learning (DRL) control discovery below.

We recognize the critical and essential nature of the initial pro-inflammatory response, the issue with VML is the extent and duration of this process, and therefore measures should be applied to biologically “close the wound.” Prior research has suggested a role for extracellular matrix (ECM) [1, 51-53], and therefore this will be incorporated into the multimodal control policy.

In terms of therapeutically addressing the late phase and attempting to modulate collagen degradation and redeposition, we recognize that the challenge here will be how to degrade Collagen III without re-activating pro-inflammatory pathways, as that is a known function of ECM-degrading enzymes such as matrix metalloproteinases (MMPs). Therefore, we deferred the investigation of this target at this phase of our work.

The following sections describe the results of the control discovery investigations, carried out in a modular fashion such that the necessity of each component can be demonstrated.

#### 2.3.2 Initial Multimodal control using mediator manipulation

Our developed approach to discovering novel multi-modal and adaptive control policies to affect cellular-molecular processes towards a specified clinical goal involves simulation-based Deep Reinforcement Learning (DRL) [54]; the specifics of this approach are discussed in the Methods and are explained at length in Refs [55, 56]. The process of simulation-based DRL is also consistent with the NASEM Report guidance on DTs, and the requirements of this approach can be identified in our specified design process for a DT [25], and which was applied to the WEABM.

As with our prior work on simulation-based DRL to cure sepsis the initial proof-of-concept implementation of DRL training on the WEABM used the most extensive sense/observation and actuate/control spaces in terms of mediator state of the wound and potential mediators for manipulation: every mediator could be observed in the 15-minute step interval of the WEABM, and every mediator could be augmented or inhibited at this interval as well. This allows us to determine the full extent of controllability given an initial representation of the potential control space based on the representation level of the WEABM; the inability to reach a desired end-state would suggest the need to expand the potential control space. With the initial maximal observation and action space the DRL training returned the top 10 performing control policies seen in Figure 10.

**Figure 10:**
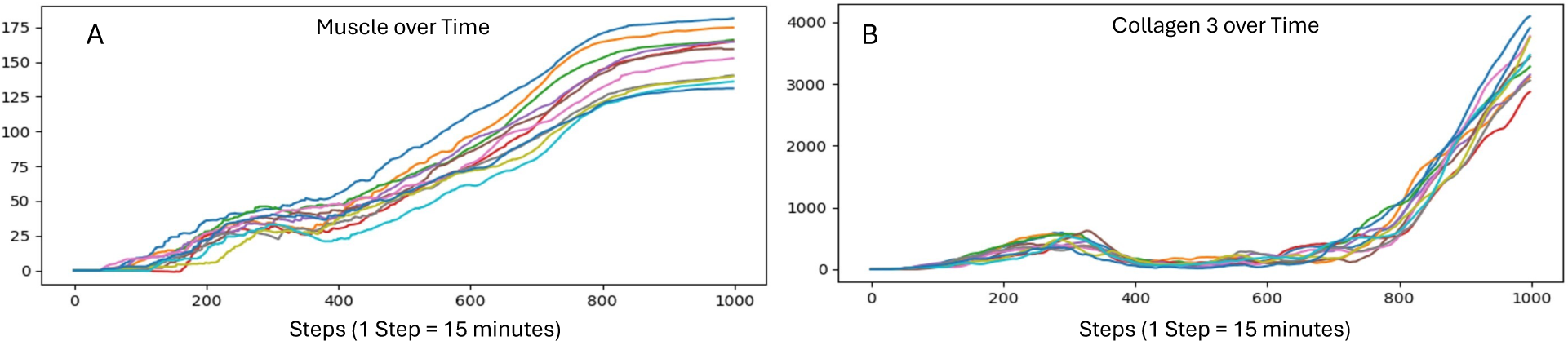
Full Control and Observation Space Top 10 DRL Policy Performance for 1000 steps = 10.4 days. Panel A shows the performance of the top 10 policies in terms of regrowth of muscle. Note that muscle regrowth is not seen at all in VML in nature, and therefore any positive curve represents a qualitative change in wound healing dynamics. Panel B shows the concurrent production and deposition of Collagen 3. Note that the end state of the wound is still primarily fibrotic, as the upward inflection in Collagen 3 production/deposition leads to the plateau of muscle regrowth.

The resulting policies are shown in a heatmap of direction of action and strength of action in Figure 11.

**Figure 11:**
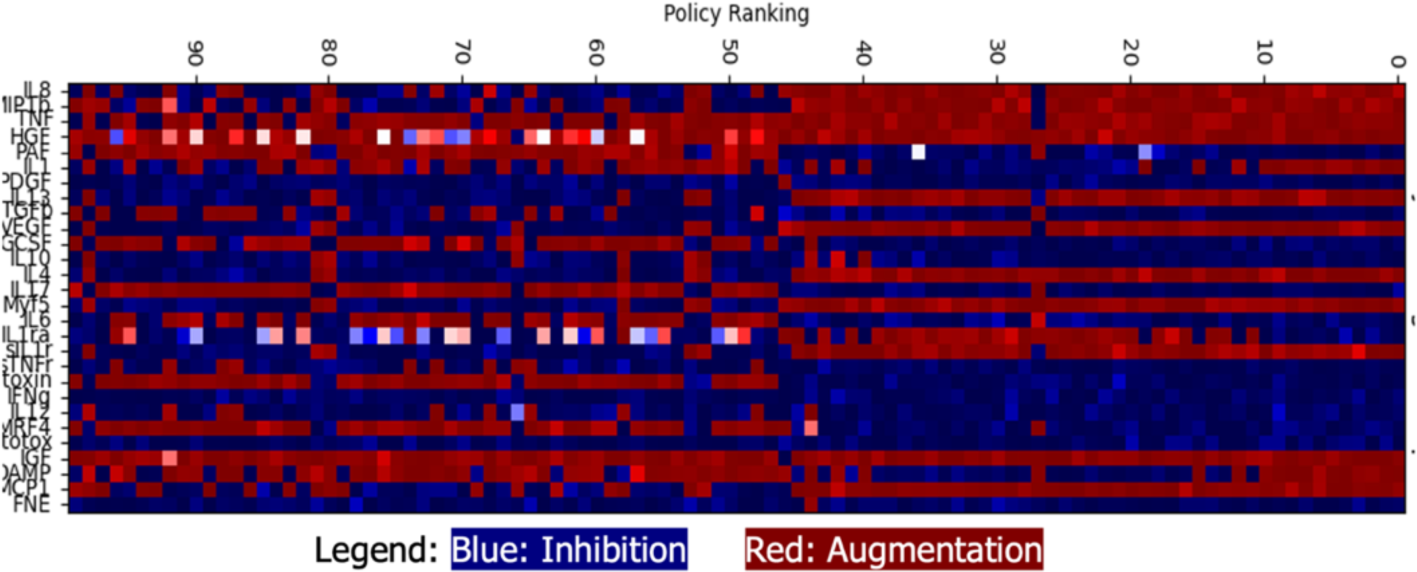
Heatmap of Policies, Top 10 policies to the Right of the figure. As noted in the Legend, Blue represents inhibition of a particular mediator whereas Red represents the augmentation of the mediator, darker shades represent higher levels of manipulation whereas lighter shades represent smaller manipulations (white = neutral).

The top 10 policies are all qualitatively similar. They are relatively static/constant through the entire 10-day course of simulated time, with actions near the maximum amplitudes of either augmentation or inhibition. The augmented mediators are: IL6, MIP1β, TNFα, HGF, IL1, IL13, VEGF, IL4, Myf5, sIL1r, IGF, a damage-associated molecular pattern (likely HMBG-1) and MCP1. The Inhibited mediators are: PAF, PDGF, TGFβ1, GCSF, IL10, IL17A, IL6, IL1ra, sTNFr, IFNγ, IL12, MRF4 and FNE.

Next, we attempted to identify a minimally sufficient observation/sense and control/actuator space by selecting canonical mediators for distinctly identified functions. The selected control targets were:

- Pro-inflammatory Mediator: TNFα
- Pro-inflammatory product: Reactive Oxygen Species (ROS)
- Anti-inflammatory Mediator: IL10
- Anti-inflammatory-Profibrosis Mediator: TGFβ1
- Pro-muscle Growth Factor: Hepatocyte Growth Factor (HGF)

In terms of the observation space, we posited that the key tissue states of interest in this phase of wound healing would be the pro-inflammatory state of the wound, which we represented by summing all the pro-inflammatory mediators, and the pro-fibrotic state of the wound, for which we selected TGFβ1 as the sensing target. We acknowledge that there might be a confounding effect with TGFβ1 as both an actuate and sense target but believe that given the central role TGFβ1 plays as a key regulator of fibrosis our initial studies would evaluate whether this dual role in DRL training would impact the resulting control policies. DRL was then performed with sensing targets of aggregated pro-inflammatory mediators and TGFβ1 and control targets of TNFα, ROS, IL10, TGFβ1 and HGF. We also increased the maximal augmentation allowed to 10-fold increase over baseline tissue levels as opposed to doubling in the prior DRL training sequence. The results of this discovered policy can be seen in Figure 12.

**Figure 12:**
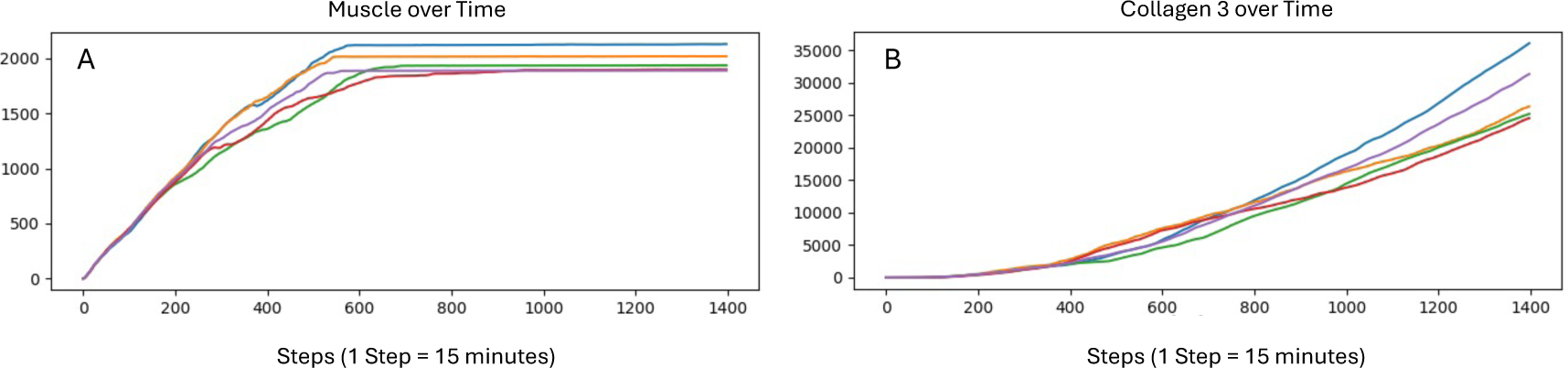
Top 10 control policies with restricted observation and action space for 1000 Steps = 10.4 days: sensing targets of aggregated pro-inflammatory mediators and TGFβ and control targets of TNFα, ROS, IL10, TGFβ and HGF. Maximal action augmentation 10x greater than employed in results seen in Figure 10. Panel A shows muscle regrowth, greater than 10-fold more muscle being produced. Panel B, however, shows a similarly increased level of Collagen 3 produced. The inflection point where muscle growth plateaus is similar to that seen in Figure 10, reinforcing the concept that the rate-limiting feature is the available damaged ends of healthy myofibrils.

These results show that with increased action magnitude there can be an increase in the amount of regrown muscle; we subsequently performed a parameter sweep of increased amplification levels that showed this effect plateaued at a 10-fold increase over baseline tissue levels. However, the plateau of muscle regrowth remains, with the significant remainder of the wound still closed with Collagen 3, which would subsequently be converted to non-functional scar tissue. With a smaller set of control targets, we are able to provide a more granular analysis of the implemented control actions and depict the time courses of the control actions of the top 10 policies in Figure 13.

**Figure 13:**
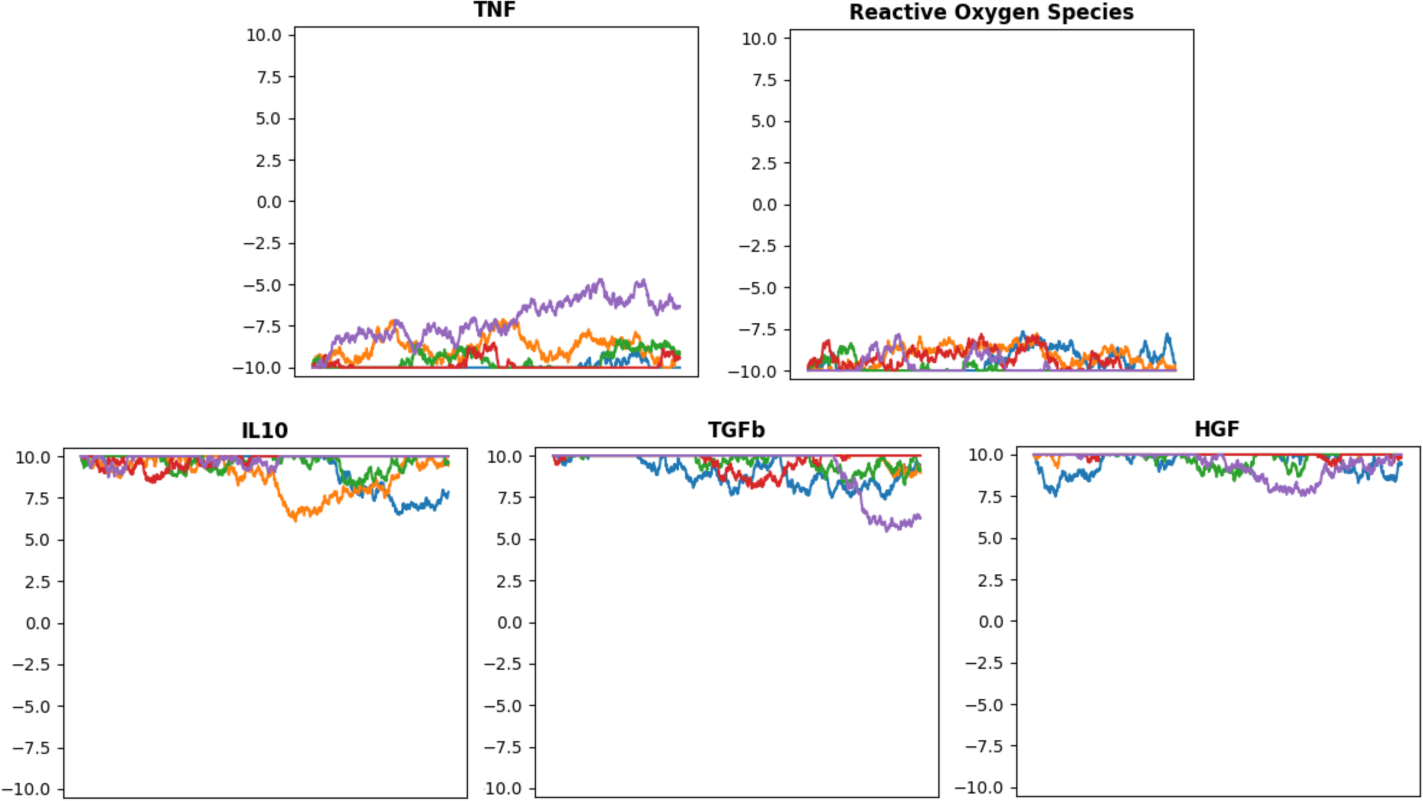
Time courses of control actions in the top 10 policies of DRL training with restricted observation and action space. The different colored lines represent different control policies arrived at via the DRL training. While there are variable levels of control from time point to time point, in general the policies implement suppression/inhibition of pro-inflammatory mediators TNFα and ROS, while augmenting IL10, TGFβ and HGF.

The control actions in the discovered control policies are relatively consistent, with inhibition of pro-inflammation and suppression of the M1 phenotype by reduction of TNFα and ROS and augmentation of IL10, TGFβ1 and HGF. Note that there is a discrepancy in the direction of control for certain mediators (specifically IL10) with the entire versus restricted control space; we interpret this as a result of restricting the control space such that it reduces the number of parallel pathways being affected. The variability seen between the depicted control actions for each individual time course is more likely to be due to variation on the specific behavioral dynamics from run arising from the stochastic nature of the WEABM. The policy is interpretable, both in terms of the desired goal of suppressing the initial pro-inflammatory phase and the need to augment myogenesis (via HGF). However, the dual effects of IL10 and TGFβ as anti-inflammatory but pro-growth/fibrosis mediators, is reflected by the enhanced degree of Collagen 3 deposition seen. Our understanding of the required effects needed to enhance muscle regrowth in VML suggests that what is needed is a more targeted anti-fibrosis controller that does not include ambiguity in terms of its targeted functions. We will return to this potential control point after we investigate the two other control modalities introduced at the beginning of this Section.

#### 2.3.2 Implementing the effect of ECM

We have noted that one of the clinical hallmarks of VML is the fact that it is an open wound that needs to be covered to avoid ongoing contamination, desiccation, and trauma. Even wound coverage with a non-bio-active dressing is itself local tissue trauma (by triggering a foreign body reaction), and therefore we consider the open nature of VML injury as inducing an ongoing source of inflammation at the tissue-atmospheric/environmental interface. This ongoing inflammatory stimulus leads to a persistence of the pro-inflammatory phase of wound healing, with consequent enhancement of the pro-fibrosis processes. Therefore, we consider an essential component of an effective control strategy to enhance muscle regrowth in VML to be a means of *biologically closing the wound* if native tissue cannot be used to accomplish this. We therefore consider the application of an artificial manufactured ECM to serve this purpose. While we recognize that there is likely a plethora of signaling effects of the ECM, these remain insufficiently characterized to the degree of detail and specificity needed to inform a mechanistic model such as the WEABM. Because of this we make the abstracting modeling decision to treat the primary effect of ECM as being the dampening of the pro-inflammatory stimulus of tissue at the surface of the wound (75% Reduction of DAMP generation at the wound surface). We then performed simulations to determine if this hypothesized mechanism is sufficient to explain experimental data that employs ECM in the canine wound. The reference model is again that which is presented in Ref [24], but now including interval application of an ECM hydrogel. The results of the MRM calibration runs are seen in Figure 14.

**Figure 14:**
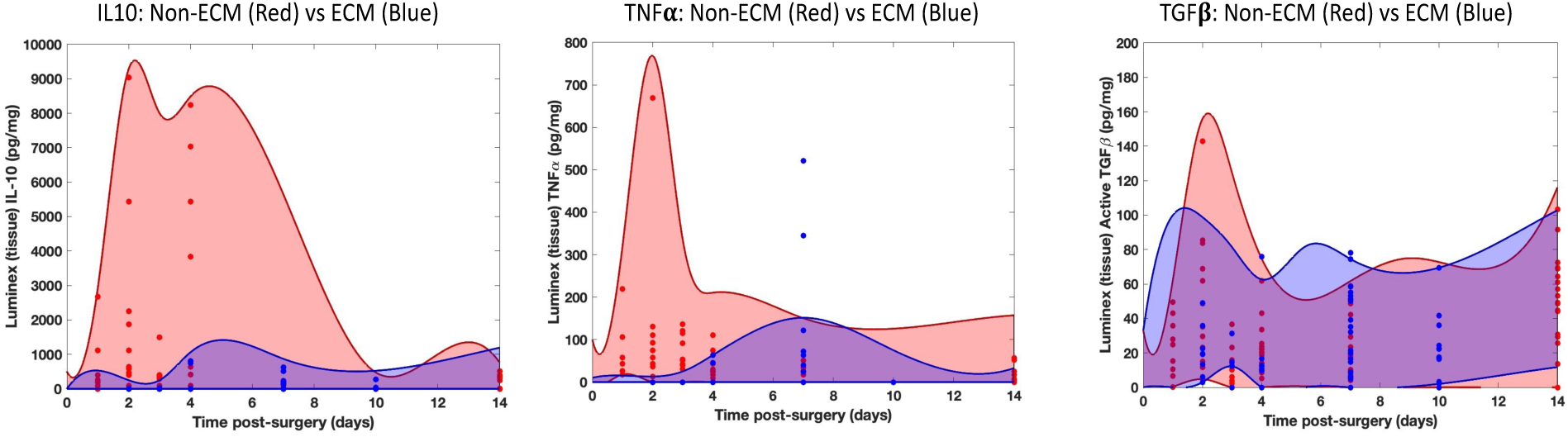
Comparison of different MRM sets fit to the baseline native healing versus with ECM application. The Red Dots are experimental data points from non-ECM animals, the Red Shaded areas are trajectory spaces of MRM configurations arrived at using the Nested Active Learning Pipeline. The Blue Dots are experimental data points from animals with ECM application within the first 10 days, the Blue Shaded areas are trajectory spaces from MRM configurations defined by the Nested Active Learning Pipeline. The ability of these two trajectory spaces to encompass their respective data while implementing the effect of ECM as 75% reduction of DAMPs at the surface of the wound demonstrates the sufficiency of that modeling assumption. Also note that there are two outlier data points in the TNFα plot; this is a similar situation to the outliers noted in Figure 7D, where out-of-bound TNFα values could be attributed to specific animals that suffered a systemic illness.

To further demonstrate the importance of ECM implemented as biologically covering the wound, we examine the efficacy of the policy discovered in Figure 13 in circumstances where the control device is applied for a limited period and then removed and standard wound care is applied during the off-control interval. These simulations involved: 1) episodic application of the control, which is assumed to include an ECM barrier to prevent atmospheric irritation of the wound surface and 2) representation of non-control intervals assumed to have standard non-biological dressings in contact with the wound surface, with consequent pro-inflammatory stimulation due to the environmental/foreign body interaction. The episode intervals were application of the control for 6 hours, followed by an off-control interval of 42 hours, for a full 48 hours of a complete episode of treatment. The questions being investigated with simulating these conditions were: 1) is a previously efficacious control policy generalizable to this different context of use, and 2) is it even possible to use DRL to train an AI control policy that would be efficacious with this context of use? The first set of simulations just tested the previously derived control policy seen in Figure 13 in a standard continuous application case, the second set of simulations tested the previously trained AI control policy in the episodic control context and the third involved implementing a DRL training program on the episodic application context. The results of these simulation experiments are seen in Figure 15.

**Figure 15:**
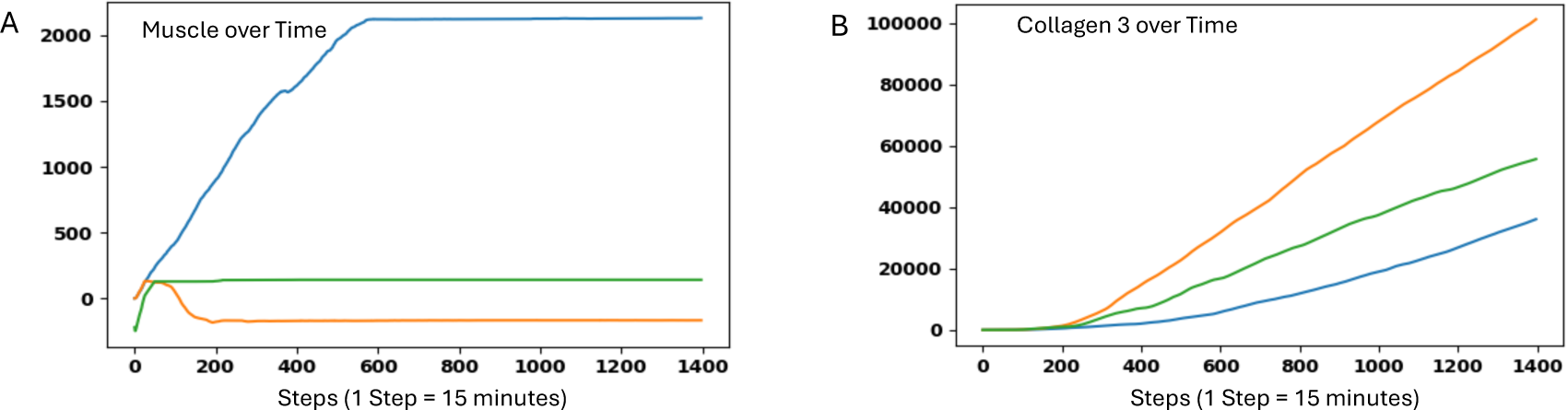
Simulations of episodic therapy for 1400 steps = 14.6 days. Panel A: Performance of Agents 1, 2 and 3 in terms of regrowth of muscle. Panel B: Performance of Agents 1, 2 and 3 in terms of generated Collagen 3. Agent 1 = AI Controller trained and reported in Figure 13. Agent 2 = AI Controller trained on episodic application of control. The blue plot shows the performance of Agent 1 in a set of WEABM simulations similar to the continuous control conditions upon which Agent 1 was trained; the performance of this control policy is equivalent to that seen in Figure 13. The orange plot shows the performance of Agent 1 in the episodic control context; in this setting the previous control policy completely fails. The green plot shows the performance of Agent 2, the AI control policy trained in the episodic context. Here there is marginal increase in muscle that quickly plateaus and becomes ineffective after about the first 48-hr control cycle.

Figure 15 shows that the previously trained AI control policy remains effective in a test condition similar to its training conditions (as expected). However, this control policy fails when applied to the episodic control context. Moreover, evening attempting to train in the episodic context does not result in an effective control policy (Agent 2). The depiction of the time series control actions for all 3 conditions is seen in Figure 16.

**Figure 16:**
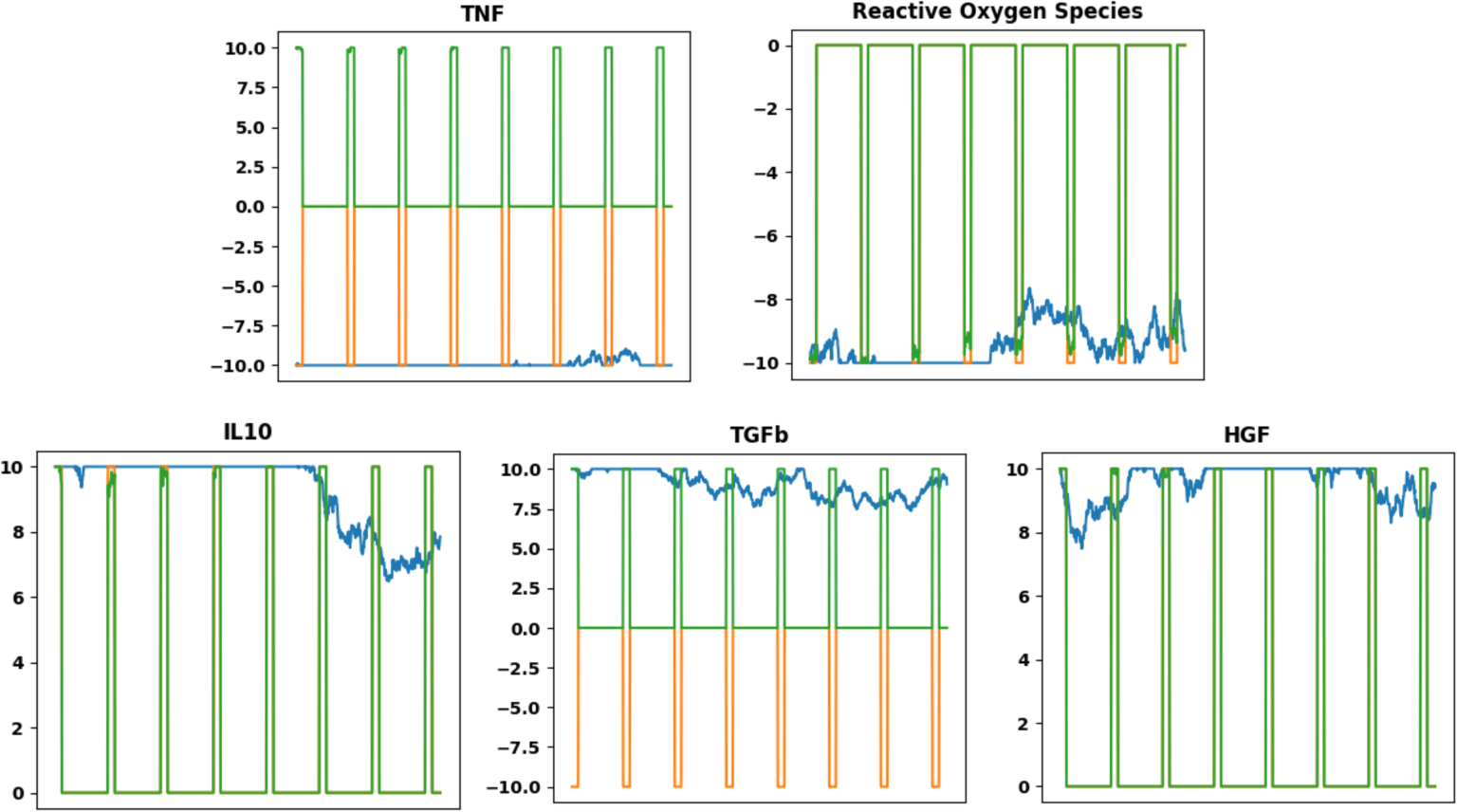
Time courses of control actions taken during episodic control application where the control is applied for 6 hours followed by a 42-hour period off-control with standard wound care. Agent 1 = AI Controller trained and reported in Figure 13. Agent 2 = AI Controller trained on episodic application of control. The blue plot shows the performance of Agent 1 in a set of WEABM simulations similar to the continuous control conditions upon which Agent 1 was trained; actions performed with this control policy are similar to those control actions in Figure 13. The orange plot shows the performance of Agent 1 in the episodic control context; in this setting the previously trained control policy generates maximal actions during the control interval that suggests its sensing of the state of the wound is continually reset after each off-control interval. The green plot shows the performance of Agent 2, the AI control policy trained in the episodic context. This policy appears to result in actions that also reset after each off-control interval that are at a maximal amplitude of intervention.

The control actions taken during the episodic application of the control with intervals off-control with standard wound care suggest that from the standpoint of the AI control agent the biology of the wound resets after each off-control interval. This is consistent with our concept of the robustness of the baseline biological imperative to close the wound, and points to the need to incorporate some means of mitigating the pro-inflammatory stimulus from the wound-atmospheric/environmental interface. We pose that ECM can serve this purpose.

#### 2.3.3 The Missing Piece: Limiting Fibroblast collagen deposition

The control modalities thus implemented and tested include:

- Manipulation of mediator milieu to attenuate pro-inflammation/M1 and foster anti-inflammation/M2 via a DRL-derived AI control policy
- Augmentation of myogenesis via HGF supplementation and electrical stimulation
- Mitigation of the pro-inflammatory stimulus from the wound-atmospheric/environmental interface from the open wound with ECM

While these control modalities can promote the regrowth of muscle in the wound (again, a phenomenon not seen in the natural case), the robustness of the fibrosis imperative still results in most of the wound filling with Collagen 3 and subsequently with non-functional scar (see Figures 10, 12 and 15). What is missing in the control space is the ability to directly mitigate the production of Collagen 3 from fibroblasts, or attenuate the differentiation of fibroblasts to myofibroblasts, which produce substantially more Collagen 3 and contract the wound. While there are known anti-fibrosis agents that have been tested in VML, such as losartan [57], the small molecule IDT1 [58] and nintedanib [59], we have elected to avoid potential off-target effects by proposing a hypothetical anti-fibrosis agent that would directly suppress the Collagen 3 production from fibroblasts as an additional control agent in a multimodal control strategy. This hypothetical compound is implemented and incorporated in the multimodal control regimen noted in the section below.

#### 2.3.5 Multimodal Control: Putting it all together

Our interpretation of the results of the different control modalities evaluated with the WEABM and our reading of the existing literature led us to conclude that there are 4 primary control areas for enhancing the regrowth of functional muscle in VML. These 4 areas are:

1. Controlling the transition from the early pro-inflammatory M1 phase to the counter-inflammation/pro-growth M2 phase in the initial response to VML injury. This we believe can be controlled at the mediator level.
2. Mitigating the pro-fibrotic aspects of the M2 pro-growth phase of the VML response by targeted suppression of fibroblast-myofibroblast function. We believe this could be accomplished by the targeted administration of known antifibrosis compounds and/or the suppression of differentiation of fibroblasts into myofibroblasts.
3. Mitigating the ongoing inflammatory stimulus from the wound-atmospheric/environmental interface that drives continued M1 stimulation and the generation of the evolutionarily beneficial wound fibrosis phenotype. We believe this can be accomplished by “closing the wound” using engineered ECM.
4. Augmenting existing myogenesis pathways, enhancing myoblast migration, fusion, and differentiation. We propose to use HGF as an endogenous compound to facilitate these mechanisms.

Underpinning these control areas are two fundamental insights:

1. There is a spatial constraint in the competition between fibrosis and myogenesis due to the limited edge/surface of damaged muscle to which any new muscle must attach.
2. The evolutionary imperative is to close the wound to limit the risk of infection. This process is fibrosis, and therefore fibrosis is a heavily favored process in the endogenous healing of VML.

Therefore, we implemented a multi-modal control strategy in the WEABM that included:

1. Mediator manipulation governed by a DRL-trained AI control policy that targeted TNFα, ROS, IL10, TGFβ1 and HGF. To demonstrate the need to include a pro-myogenesis mediator an additional control policy was generated with no HGF.
2. The addition of a hypothetical anti-fibrosis compound that selective reduces fibroblast production of collagen (and subsequent differentiation into myofibroblasts).
3. A surface covering with an ECM that dampens the pro-inflammatory stimulus from the open surface of the wound.

To evaluate the contribution of each of these components in the overall performance of a control strategy, combinations of the control modalities were examined for efficacy for muscle regrowth. The results of these simulation experiments are seen in Figure 18.

**Figure 18:**
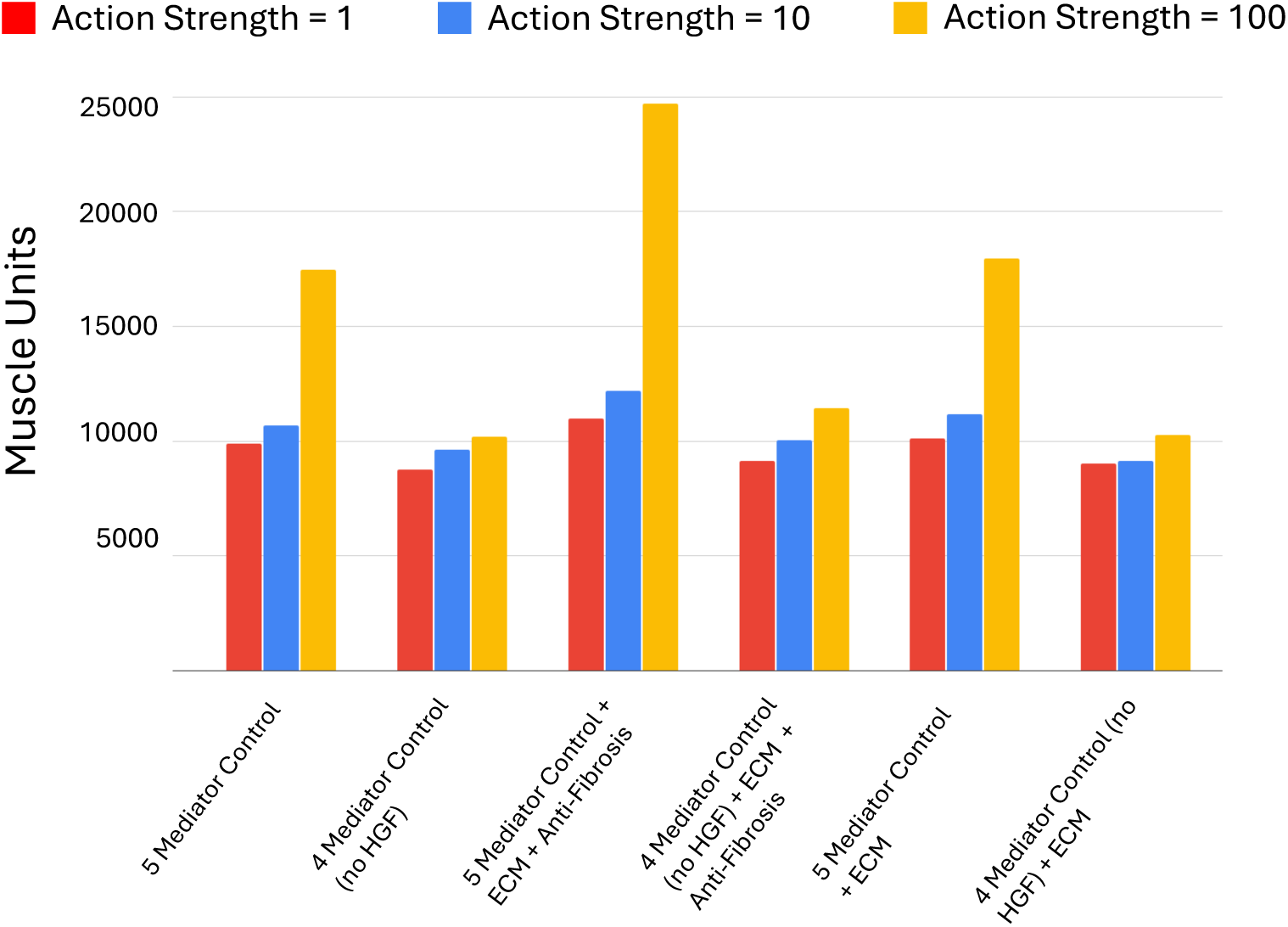
Comparison in muscle regrowth between different combinations of control strategies with 3 different levels of mediator manipulation implemented in the DRL training. In all cases the strongest level of manipulation (100-fold increase over baseline tissue mediator level) led to the greatest muscle growth.

All the treatment regimens included a DRL-trained AI control policy that governed mediator manipulation (either all 5 above noted mediators or a set of 4 minus HGF); each combination included DRL training at 3 levels of maximal mediator manipulation, 1x, 10x, or 100x over baseline tissue levels. In all cases, the 100-fold increase over baseline led to the greatest amount of muscle regrowth, demonstrating the resistance of the underlying fibrosis trajectory to alteration. These results also show the critical role of a pro-myogenesis compound (in this case HGF) in facilitating muscle regrowth. Finally, the greatest amount of muscle regrowth required the addition of an anti-fibrosis compound that directly targeted fibroblast function and/or differentiation to myofibroblasts.

## 3.0 Summary

The WEABM fits into the DT Paradigm as described in the NASEM Report in the follow ways:

- Personalization at Initialization: Uses CT topography to initialize computational model
- Incorporates an ongoing data linkage between the real- and digital-twin (sense/actuate capability)
- Personalization based on updatable forecasting
- Personalization in terms of adaptive control

Biological insights provided by the development and use of the WEABM include:

- Competition between fibrosis and myogenesis due to spatial constraints on available edges of intact myofibrils to initiate the myoblast differentiation process.
- Need to biologically “close” the wound from atmospheric/environmental exposure, which represents an ongoing inflammatory stimulus and “resets” the disease process into the early M1 phase, also dynamics seen with chronic non-healing wounds.
- Selective, multimodal and adaptive control needed, inherent to this is the DT concept of ongoing sensing/data link between the virtual and real world.

Though we can generally characterize the dynamic phases of wound healing in VML, a primarily pro-inflammatory response in the first 3 days, with a transition to a predominance of anti-inflammatory functions from that time to about 7-10 days, followed by an initial acceleration of collagen production and early fibrosis from approximately 7-14 days, followed by collagen remodeling and the eventual formation of fibrotic scar, we also recognize that there is considerable heterogeneity both between individuals and within an individual’s wound. Because of this, the ability of the control AI to adapt its recommended multi-mediator intervention to the state of the wound is a key aspect of being able to override the biological imperative leading to fibrosis. Based on the simulation experiments and DRL-based control discovery with the WEABM we would propose the following design specifications for a device to be applied to the surface of the wound:

- Sensing Capabilities:
  - Characterization of Pro-inflammatory State: We would suggest IL6 as a sense target given its well-known role as an aggregated marker of inflammation.
  - Characterization of Anti-inflammatory and Pro-growth State: We would suggest TGFβ as a marker for the translation between beneficial anti-inflammatory conditions and potentially pro-fibrosis conditions.
- Control Capabilities:
  - Variable delivery of the following mediators (minimal set) at levels up to 100-fold increase over maximum endogenous tissue levels or near complete inhibition via direct antagonists (either chemical agents or monoclonal antibodies):
    - Pro-inflammatory mediator: TNFα
    - Pro-inflammatory byproduct: ROS
    - Early Anti-inflammatory mediator: IL10
    - Transitional Anti-inflammatory/pro-fibrosis mediator: TGFβ1
    - Pro-myogenesis mediator: HGF
    - Anti-fibroblast/myofibroblast Mediator: Hypothetical, though could use IDT1 as a small molecule
  - An ECM-like interface with the wound surface

We propose that a device with these specifications can be a first-generation device that would be refined through its use and testing. The currently listed set of target mediators are readily modifiable based on the current version of the WEABM, and an expanded WEABM could introduce additional control targets (i.e., a more detailed representation of fibroblast function and differentiation into myofibroblasts).

As with all computational models the WEABM includes selective abstractions and is inherently incomplete. We can address the latter issue via our Model Calibration method (see Methods); the former can be refined should more detailed information become available. We have also attempted to avoid off-target effects of control compounds given the dynamic and adaptive nature of the needed control policies, and therefore have focused on delivering specific mediators and their direct antagonists. We recognize that we have heavily abstracted the potential biological interactions of the ECM, but this is due to the general lack of direct mechanistic signaling information regarding its interaction with specific cell types and regarding specific pathways of interest. Our current abstraction where the ECM serves primarily as a “inflammation sink” to mitigate the open nature of clinical VMLs has been sufficient to explain all the data generated in the associated reference canine model. A significant future aspect of clinical VMLs that will have to be addressed is the impact of microbial interactions with the wound, with attendant issues regarding how to suppress microbial virulence and pathogenicity by additional manipulation of the same host response that is governing the process of healing.

We cannot emphasize enough that the current version of the WEABM is vastly under-constrained with respect to the set of non-falsifiable configurations, and as such the proposed development and refinement process of the VML DT cyberphysical system requires the application and testing of the device *in situ* to generate the necessary iterative refinement loop. We assert that solving the problem of VML will require such an approach and hope that the introduction of this VML DT can both spur device development and provide a beneficial example of the application of the Medical DT paradigm to solving a complex and significant clinical problem.

## 4.0 Methods

### 4.1 Model Calibration

The level of abstraction implemented in this model (and our previous work) is informed by the rationale above, specifically, that the model is a representation of existing biological knowledge (noting that this does not preclude the model from positing biological features that should be investigated experimentally). This type of knowledge is typically of the form, “*cytokine x* upregulates *cytokine y*” or “*cytokine z* is necessary for this cell to perform a specific function.” This is distinct from approaches that claim to be *ab initio* or from “first principles,” as these approaches are not tractable to model organ/tissue systems at the desired level of resolution, of which a primary focus is representing biological/genetic/epigenetic heterogeneity.

Variation in biomedical data arises from two primary sources: stochastic responses to identical perturbations and genetic heterogeneity. We attempt to capture both of these effects in our modeling through both the stochastic replication of simulations with identical parameterizations and through the discovery of the set of all parameterizations that, when instantiated into the knowledge/rule structure of the model, cannot be invalidated by the available data (or some other established knowledge).[50, 60] This gives us an ensemble of putative model parameterizations that fit existing data, recognizing (and potentially hoping) that future experiments may invalidate some members of the ensemble. The result is not a model that is complete (as representing every possible biological interaction is intractable), but a model that is sufficient to explain biomedical phenomena (i.e., how the system works) in the context that there exists unavoidable epistemic incompleteness/uncertainty.

We recognize that the NASEM Report placed considerable emphasis on the importance of VVUQ for DT technology; as noted above VVUQ is the focus of NASEM Report Conclusions 2-2 through 2-4 and Recommendation 2. Specifically, there is a call for new approaches towards what may constitute VVUQ for digital twins based on their specified “fit for purpose.” We support this conclusion and have asserted that there are particular properties associated with attempting to computationally represent biomedical systems in a multi-scale fashion, specifically being able to cross cellular-molecular biology to whole patient/clinical phenotypes. We identify two key facts about biological systems that distinguish them from industrial/engineered systems and present intractable challenges to attempts to translate approaches known to be effective in many physical sciences and engineered systems:

- Biological systems are faced with perpetual epistemic uncertainty.
- Biological data, particularly time series data needed to establish system trajectories, will be perpetually sparse with respect to the range of possible data configurations.

We have presented an extensive discussion regarding the implications of these two features of biological systems in Ref [49]; we present a condensed discussion of herein. We pose that the nature of the mapping between computational representations and real-world biology is a mapping between two systems with unquantifiable uncertainty. There is uncertainty (perpetual and unquantifiable) in both the specification of the computational model (perpetual epistemic uncertainty) and the probability distributions of the real-world data (data sparsity and biological heterogeneity). Traditional approaches to validation of computational models are fundamentally incapable of dealing with this type of uncertainty mapping. However, we recognize that with a goal of defining robust control one needs to avoid failure of the proposed control in as broad a context as possible. Therefore, in cases where there are perpetual uncertainties in both the model and the data it is therefore essential to maximize the conservation of information in the how these systems are related to each other, e.g., “communicate” with each other. This insight leads to casting the Validation and Uncertainty description issue for medical DTs in the context of Shannon Information (from Information Theory) and the Maximal Entropy Principle (MEP) from Statistical Mechanics. These are analogous principles where the goal is to account for the uncertainty inherent in systems and their relationship/communication to each other by maintaining the largest set of possible configurations that provide concordance between those systems; i.e., Information as per Information Theory and Entropy as per Statistical Mechanics. The maximizing entropy/information content provides the broadest non-falsifiable set of mechanistic configurations capturing biological and clinical heterogeneity and upon which control can be discovered. Since, however, any putative control policy requires identification of ostensible control points, and any mechanism-based model must have a minimal set of represented components and interactions, we recognize that the discovery of robust control must use a mechanism-based structure that, while acknowledged to be incomplete, needs to be able to be expanded upon to meet the MEP. The task, then, is to design an approach that maximizes the information content given an initial knowledge structure.

Towards this end we have developed a mathematical object called the Model Rule Matrix, or MRM [48]. The MRM is a matrix that consists of columns that list all the entities/molecules/mediators chosen to be included in a simulation model and row that list all the rules representing biological functions/interactions chosen to be in the simulation model. The numerical values of each matrix element denote the strength and direction (inhibition or augmentation) of the contribution of the entity (column) to the functional rule (row). See Figure 2 for a depiction of the components of an MRM. The lack of inclusion of a particular entity in a particular rule is represented by a “0” for the corresponding matrix element in the initial/base MRM; this represents the choices made by the modeler regarding what to include in the computational model, which in turn is informed by what data types are available to link the model to the real-world and provides a set of putative control points.

As a result, the base MRM is sparse with most of the possible interactions not explicitly represented in the computational model. However, the selection of what is represented in the model represents a bias with respect to the full biological complexity (though an unavoidable one) and that there are numerous other potential/inevitable contributions from biological components/molecules/pathways/genes that influence the behavior of the explicitly represented model components. Thus the “0” elements in the base MRM represents a representation of a “latent” space of uncharacterized and unspecified interactions that are nonetheless known to be present in some form at some degree. We assert that it is the gap between the explicitly represented structure of a computational model and the recognized additional potential interactions (by whatever degree of connectivity) that contributes to a model to capture the richness (manifested as behavioral heterogeneity) in the real biological system. It is in this fashion that the MRM utilizes the Maximal Entropy Principle: the potential information content of the model is enriched by the representation of connections that are unknown or electively omitted that become necessary to represent the full heterogeneity of a data set. The enriched MRM is then capable of representing a genetically, epigenetically, and functionally diverse cohort of in silico patients able to represent a range of heterogeneous experimental or clinical data. The process of evolving enriched MRMs involves the application of a ML pipeline that employs both Genetic Algorithms (GAs) [61-64] and Active Learning (AL) [65-69]. to a simulation model constructed such that the coefficients of the rules in the simulation model can be considered strengths of interactions of their associated variables (and is therefore able to be represented by an MRM) and a data set that manifests a large degree of variability. The ML pipeline identifies the set of MRMs (e.g., set of possible additional connectivity configurations) that are able to encompass the range of variability present in the data set. A more detailed description of the ML pipeline can be found at Ref [48, 50].

The large degree of variability in the data is desirable because the goal is to represent the widest range of biological behavior as possible; having such a target data set expands the descriptive comprehensiveness of an ensemble of MRMs. As opposed to classical parameter fitting, which seeks to find a minimal set (or single) optimal parameter configuration(s), this process does the exact opposite by identifying a very large set of non-falsifiable configurations that has the capability of generalizing beyond the training data set and reducing the risk of brittle, over-fitted models. Time series data is particularly desirable, because the requirement of characterizing of behavioral trajectories further constrains the set of non-falsifiable MRMs (because the knowledge-based form of the rules limits their possible behaviors).

Note that this approach addresses a series of Findings and Conclusions in the NASEM Report concerning fidelity of models, multiscale modeling and model refinement. We consider all these issues related to and inherent in the establishment of novel validity and uncertainty quantification methods such as presented above.

### 4.2 Control Discovery of Complex Systems: Beyond classical control using Deep Reinforcement Learning

While there are numerous classical control approaches to complex dynamical systems, we have previously suggested that for complex biological systems that are represented by agent-based models, such control approaches are limited in their applicability [70]. Instead, we have suggested that Deep Reinforcement Learning (DRL), a form of training ANNs that utilizes time series/outcome data to identify combinations of actions that can alter/change the behavior/trajectories of a system [54], can be applied to simulation models to discover robust control policies to cure sepsis [55, 56]. Our proposed approach is analogous to the successful application of DRL to playing and winning games [71-73]. We term this group of applications of DRL as simulation-based DRL; this describes the use of simulation-generated training data without a pre-defined structural model for the DRL agent. We term this approach *simulation-based DRL*. A key feature of this approach is that since it relies on actions taken on a simulation model there is transparency in the mechanistic state transitions of the simulated system and this allows for a degree of interpretability in the putative control action that is not possible in “standard” data-centric ML and AI systems. This feature has potential significant impact when it comes time to evaluate such a system from a regulatory standpoint on a path to clinical implementation. The details of the process of training a DRL AI can be seen in Ref [56], and are summarized here.

We utilize a Deep Deterministic Policy Gradient (DDPG) [54] to train an AI to cure sepsis. DDPG is a reinforcement learning (RL) algorithm that uses off-policy data and the Bellman equation (Equation 1) to learn a value function, or Q-function, to determine the most valuable action to take given any state of the simulation (obtained by some means of observation, e.g. a sensor in the biomedical case).

**Equation 1:**
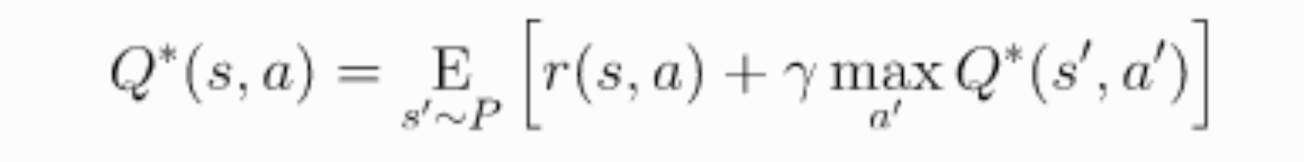
The Bellman Equation. Value Q is a function of the current state and action (s, a), and is equal to the reward r from the current state and chosen action (s, a) summed with the discounted value of the next state (discount factor = γ) and action (s’, a’) where the next state is sampled from a probability distribution (s’ ∼ P).

The Q-function is discovered through the execution of numerous simulations of the target system in which the AI agent uses trial and error to optimize the Q-function based on observed rewards from chosen actions. The basis of this process is a discrete version of Q-learning [74], where the next action taken by the AI is chosen from a set of discrete actions with the best action for the current state is the one that returns the highest value from the Q-function. Q-learning is an off-policy algorithm, which means that in the training phase the AI agent is able to choose actions not chosen by the Q-function, allowing the AI agent to explore and potentially discover actions that can lead to a greater reward than an already discovered policy. Q-learning has been demonstrated to be very effective at solving complex control problems in discrete space [54].

The full observation and action space for the WEABM consists of every mediator represented in the WEABM. This was used for the initial DRL shown in Figure 10. However, in terms of practical feasibility the results of the initial control policy were examined, and a subsequent restricted observation and control space were defined consisting of TNFα, ROS, IL10, TGFβ and HGF and eventually a hypothetical anti-fibrotic agent. Each training episode included ∼ 10,000 simulations of up to 14 days of simulated time (= 1400 steps) distributed over 500 processors; these episodes took ∼ 30 hours to execute. Specific simulation experiments are described in the Results.

## Acknowledgements

This work was supported in part by the National Institutes of Health Award UO1EB025825. This research is sponsored by the Defense Advanced Research Projects Agency (DARPA) through Cooperative Agreement D20AC00002 awarded by the U.S. Department of the Interior (DOI), Interior Business Center. The content of the information does not necessarily reflect the position or the policy of the Government, and no official endorsement should be inferred.

Special Acknowledgements to Dr. Stephen F Badylak and Scott Johnson from the University of Pittsburgh McGowan Institute of Regenerative Medicine for their work on the experimental canine model reported in Ref [24] and which provided the calibrating data for the WEABM presented herein.

## Appendix A: Facts and References for WEABM

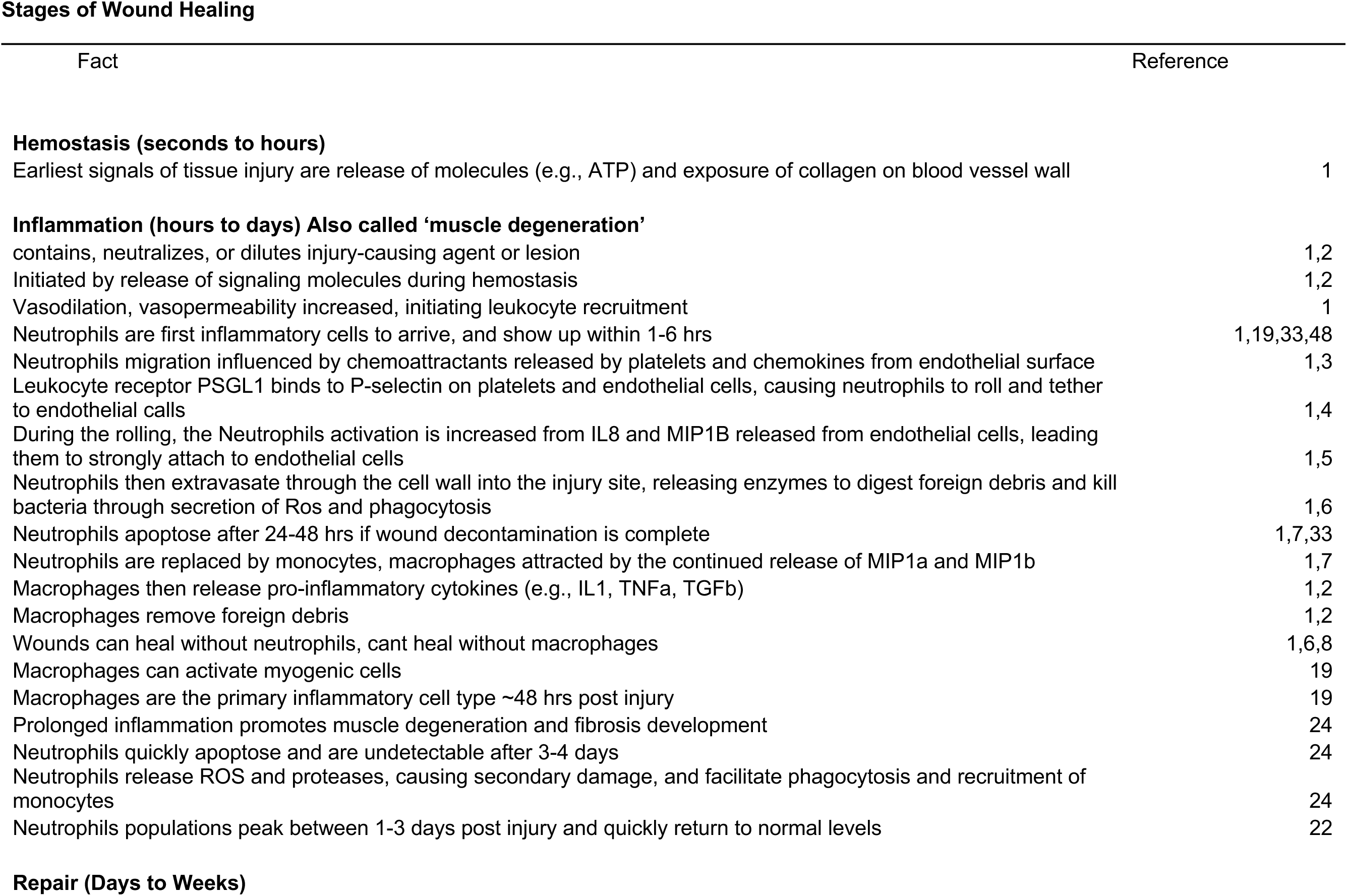

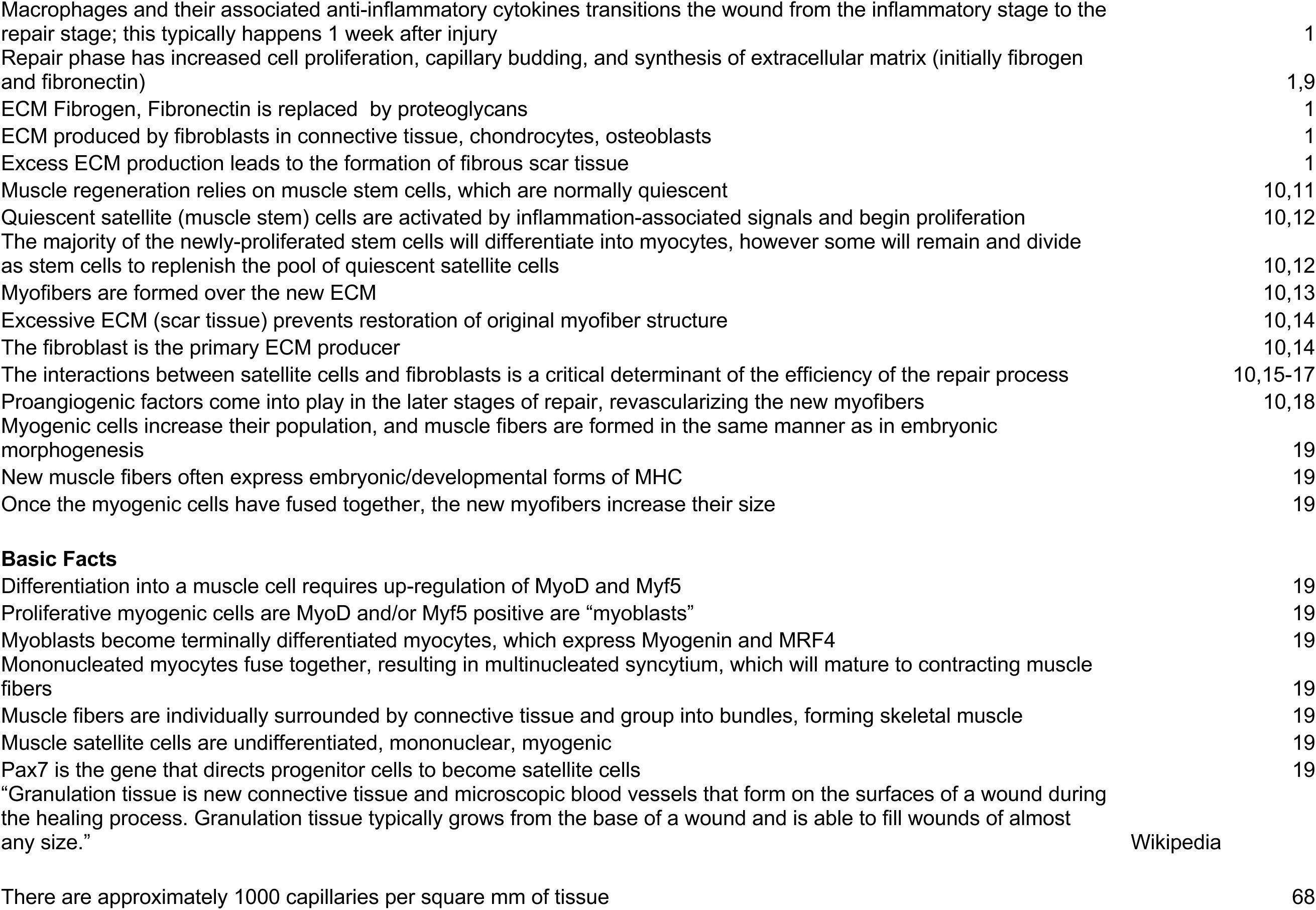

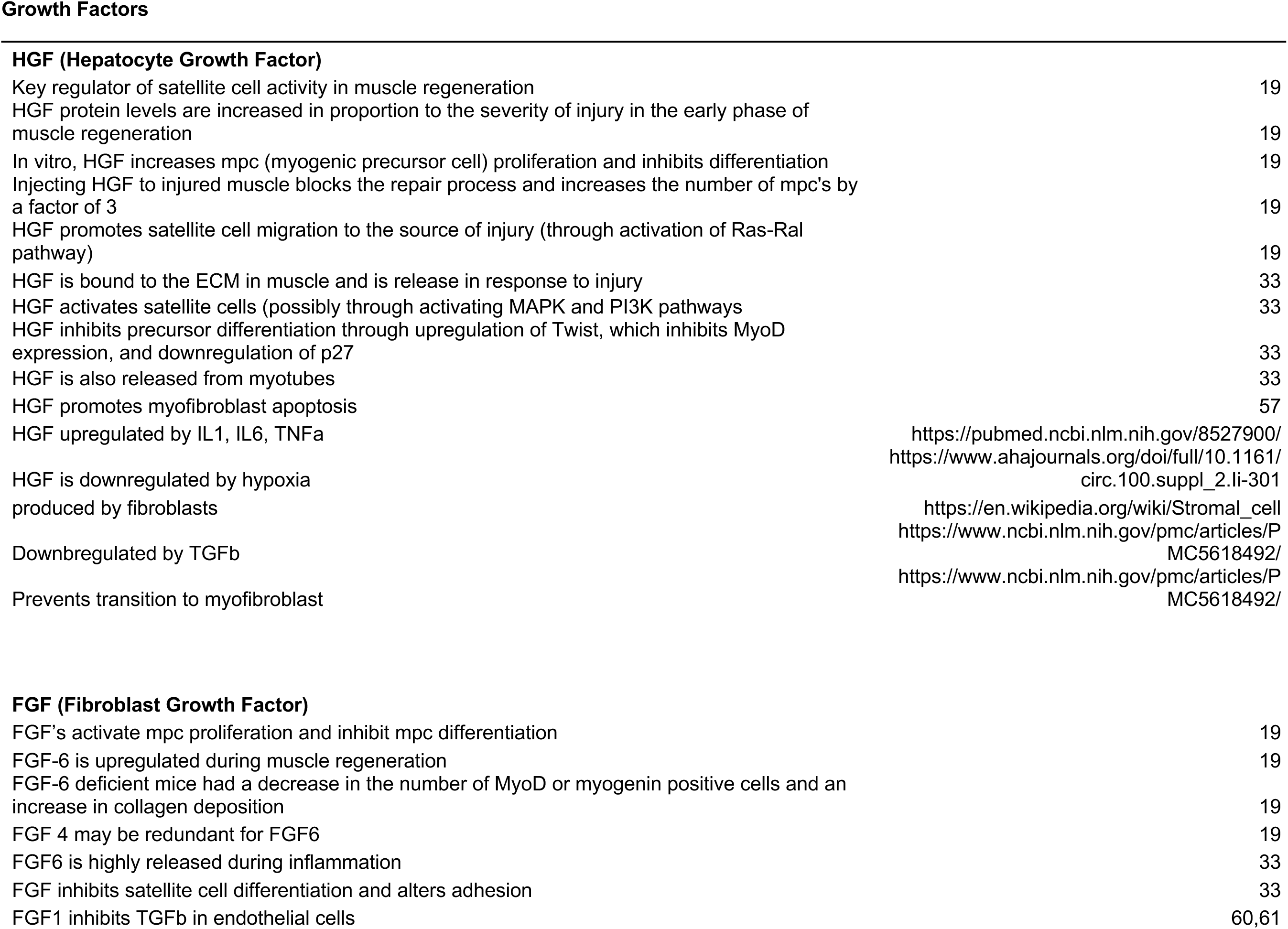

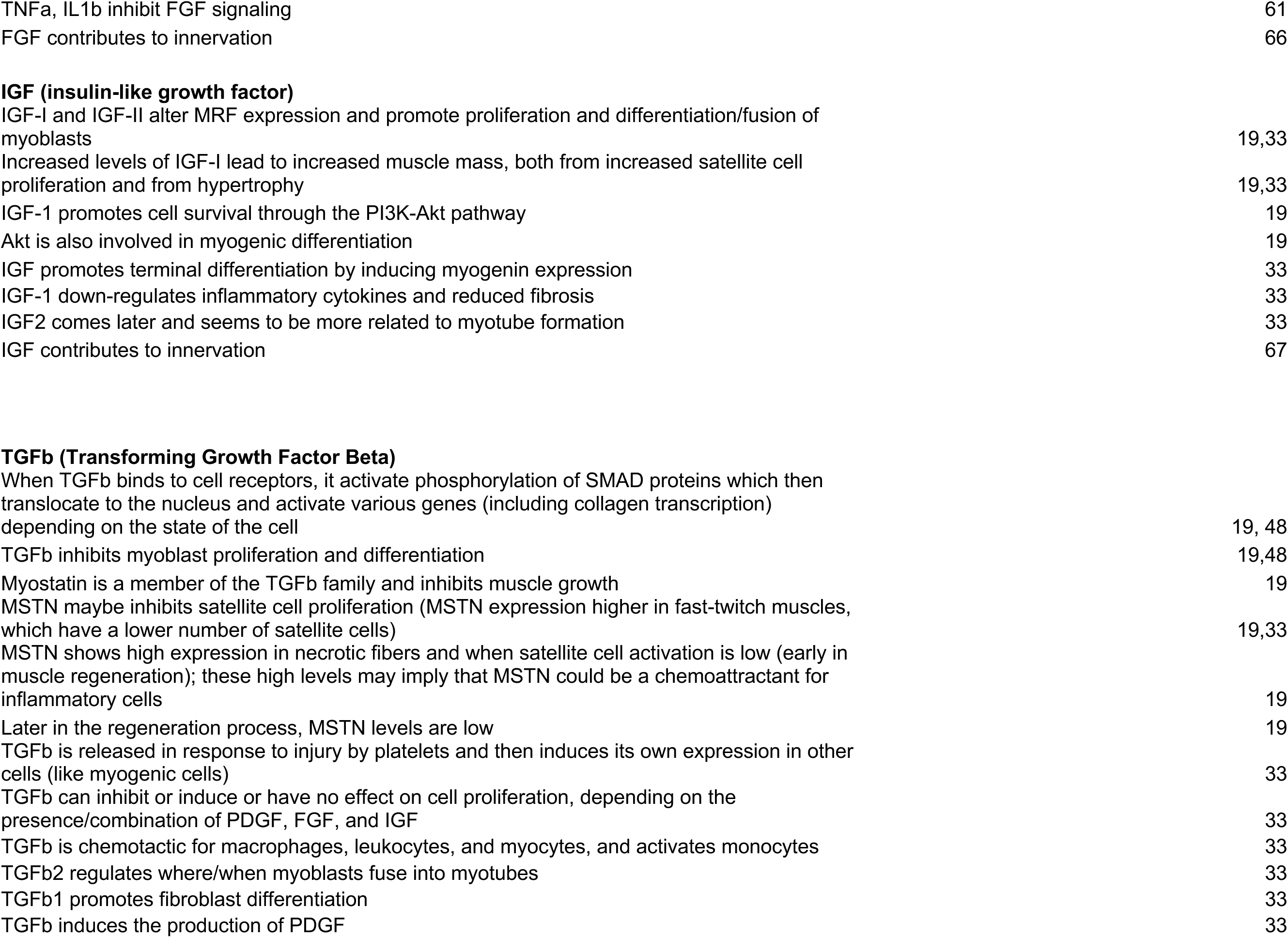

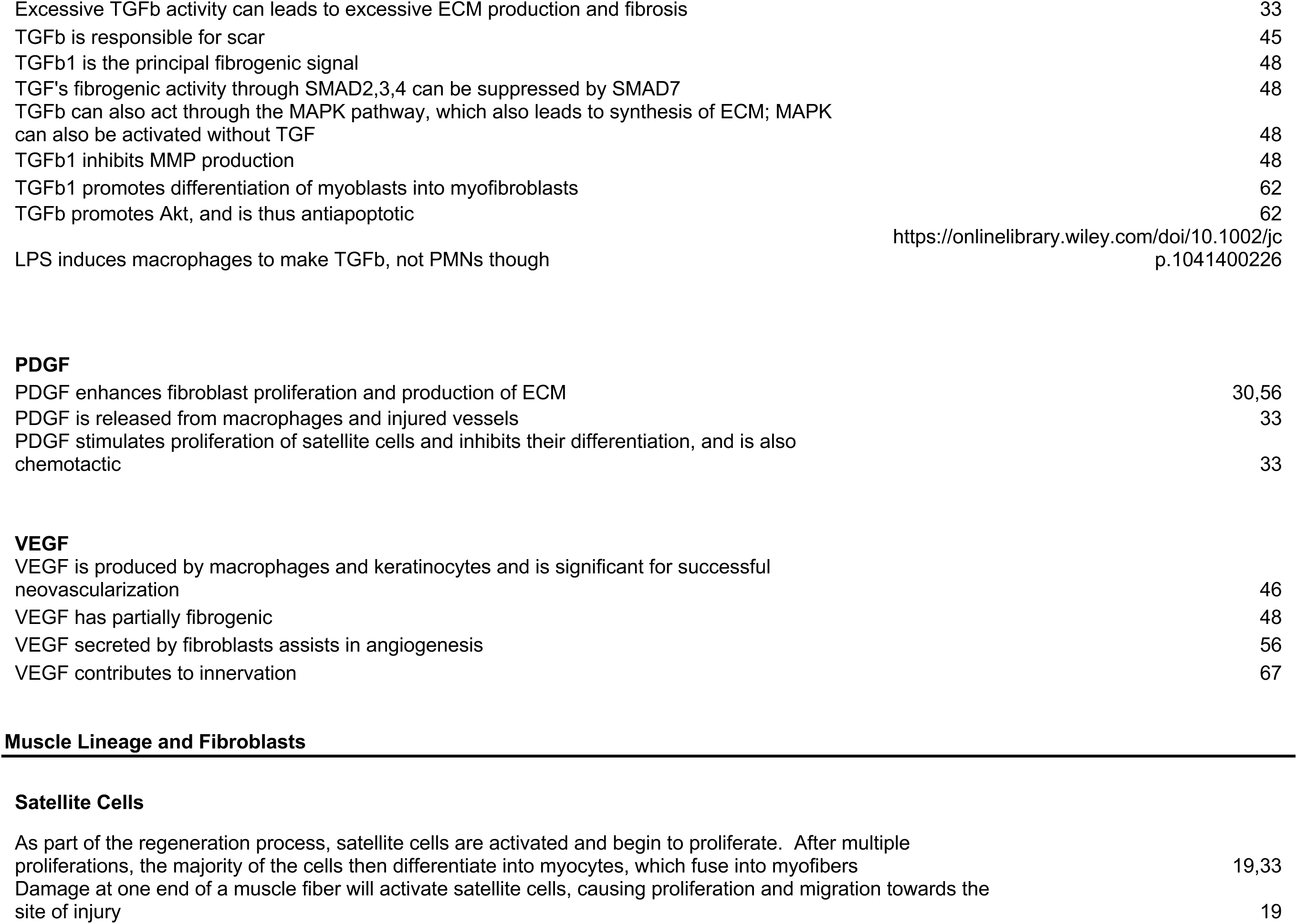

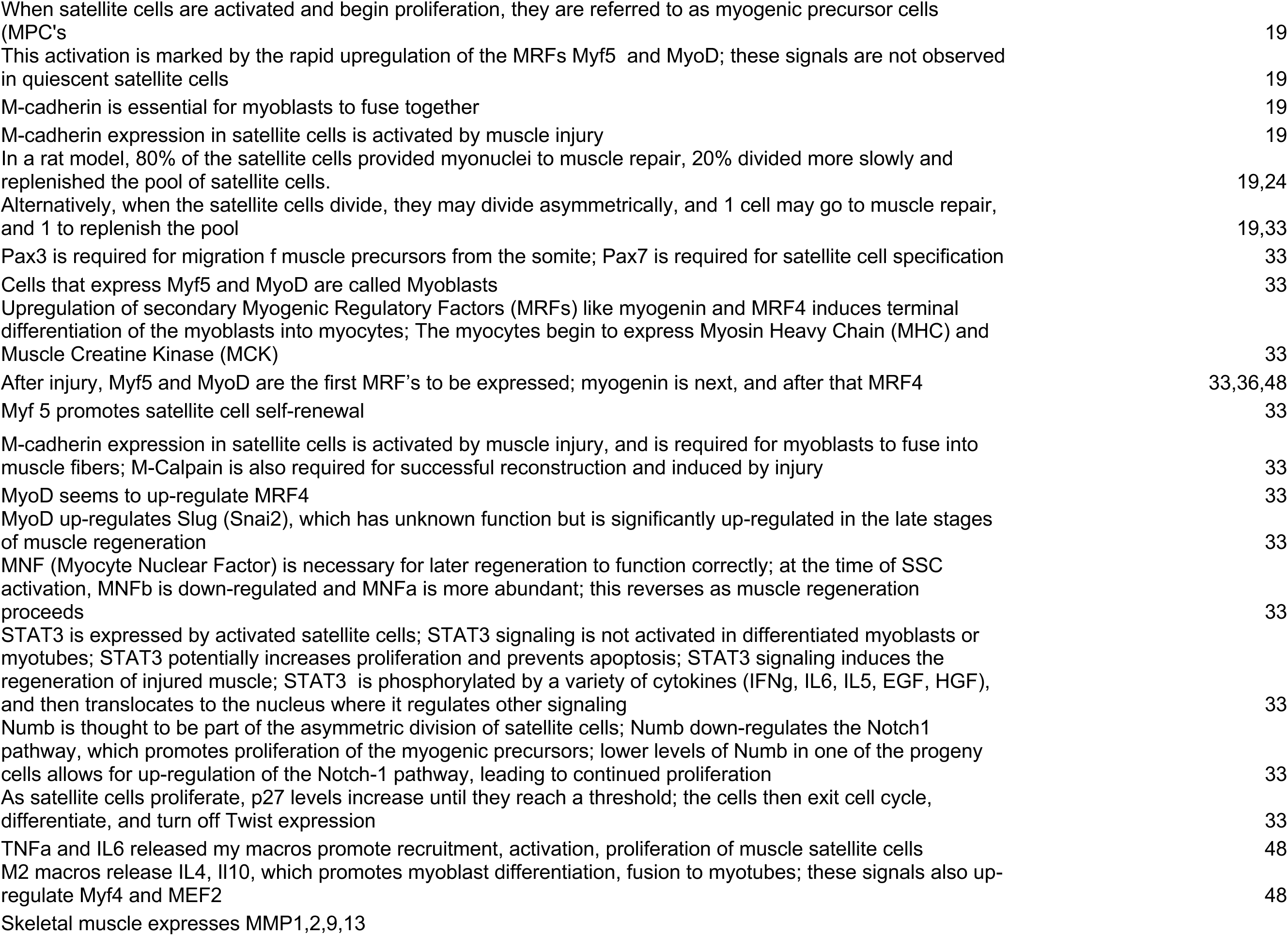

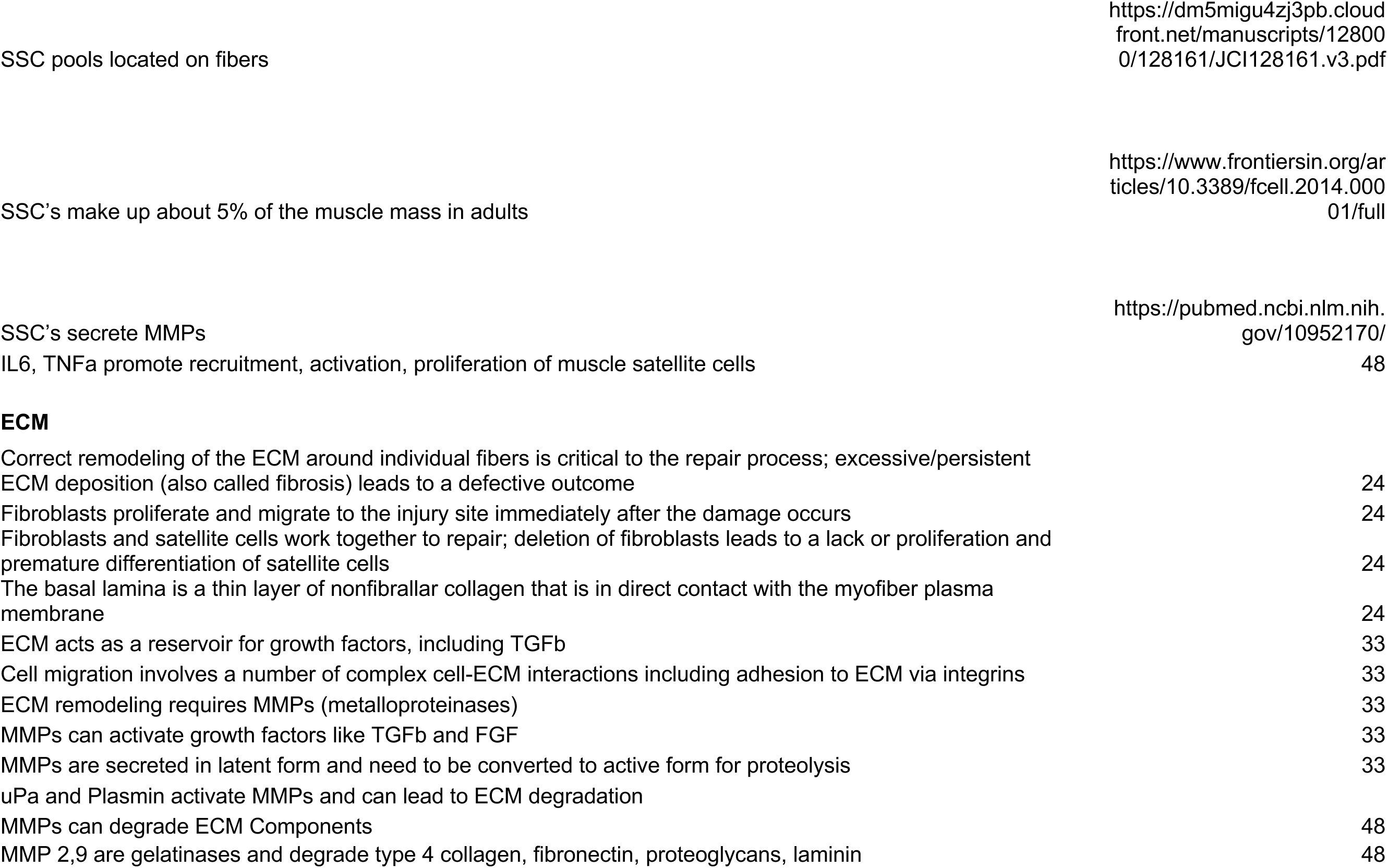

|  |  |
| --- | --- |
| MMP1,13 degrade the 1,3 collagen | 48 |
| MMP9 is up-regulated by NfKB, MAPK, STAT3 | 51 |
| IL1b, TGFb up-regulate MMP2,9 | 52,53 |
| In the synthesis of ECM process, collagen type 3 (major component of granulation tissue) is replaced by collagen type 1 | 54 |
| Type 1 collagen is associated with scar; normal tissue is a mix of type 1 and type 3, scar is mostly type 1 | 58 |
| TNFa or IL1b induced MMP1 secretion is mediated by MEK/ERK, so a MEK/ERK inhibitor will significantly inhibit the MMP secretion when induced by pro-inflammatory signals | 71 |
| **Fibroblasts** |  |
| TGFb promotes fibroblast proliferation, ECM production | 31,56 |
| ECM deposition occurs faster than myogenesis; excess ECM leads to scar tissue | 31,48 |
| TGFb can prevent Fibroblast apoptosis (triggered by TNF) | 32 |
| Fibrogenesis occurs the same way, regardless of how injury occurred (toxic, trauma, etc) | 33 |
| Ablation of fibroblast population leads to premature satellite cell differentiation and depletion of the satellite cell pool, and ultimately, smaller myofibers (maybe through TCF4) | 39 |
| TGFb1 causes fibroblasts to transform into myofibroblasts, which express alpha-smooth-muscle-actin (aSMA) | 48 |
| Fibronectin and Tenascin-C are produced first, followed by Collagen type III and collagen type I | 48 |
| For injuries that have persistent inflammation (i.e., VML), myofibroblasts do not undergo apoptosis, and remain in granulation tissue; the sustained presence of immune cells in this tissue causes the release of fibrogenic cytokines like TGFb1; this causes myofibroblasts to continue proliferation and production of ECM, which ultimately leads to scar | 48 |
| Fibroblasts secrete type 1 and 3 collagen | 55 |
| Fibroblasts can produce both MMPs and their inhibitors (TIMPs) | 56 |
| The fibroblast to myofibroblast transition enables the contraction of the ECM to seal open wounds | 56 |
| Fibroblasts are stimulated to differentiate into myofibroblasts from pro-inflammatory signals | 56 |
| Fibroblasts secrete TGFb1, IL1b, assisting in the activation/migration of macrophages | 56 |
| Fibroblasts can contribute to chronic inflammation, promoting myoblast differentiation and thus, fibrosis | 56 |
| Fibroblasts secrete VEGF, which is important for angiogenesis | 56 |
| IL13 stimulates (in fibroblasts) TGFb production, collagen production, MMP expression, fibroblast proliferation, myofibroblast differentiation | 56 |
| TGFb induces ROS production in fibroblasts | 69 |
| IL1 and IFNg reduce aSMA formation (preventing differentiation) | 69 |
| When fibroblasts are exposed to media from muscle cells which are treated with SMAD3 and TGFb, they preferentially express collagen3 over collagen 1 | 70 |

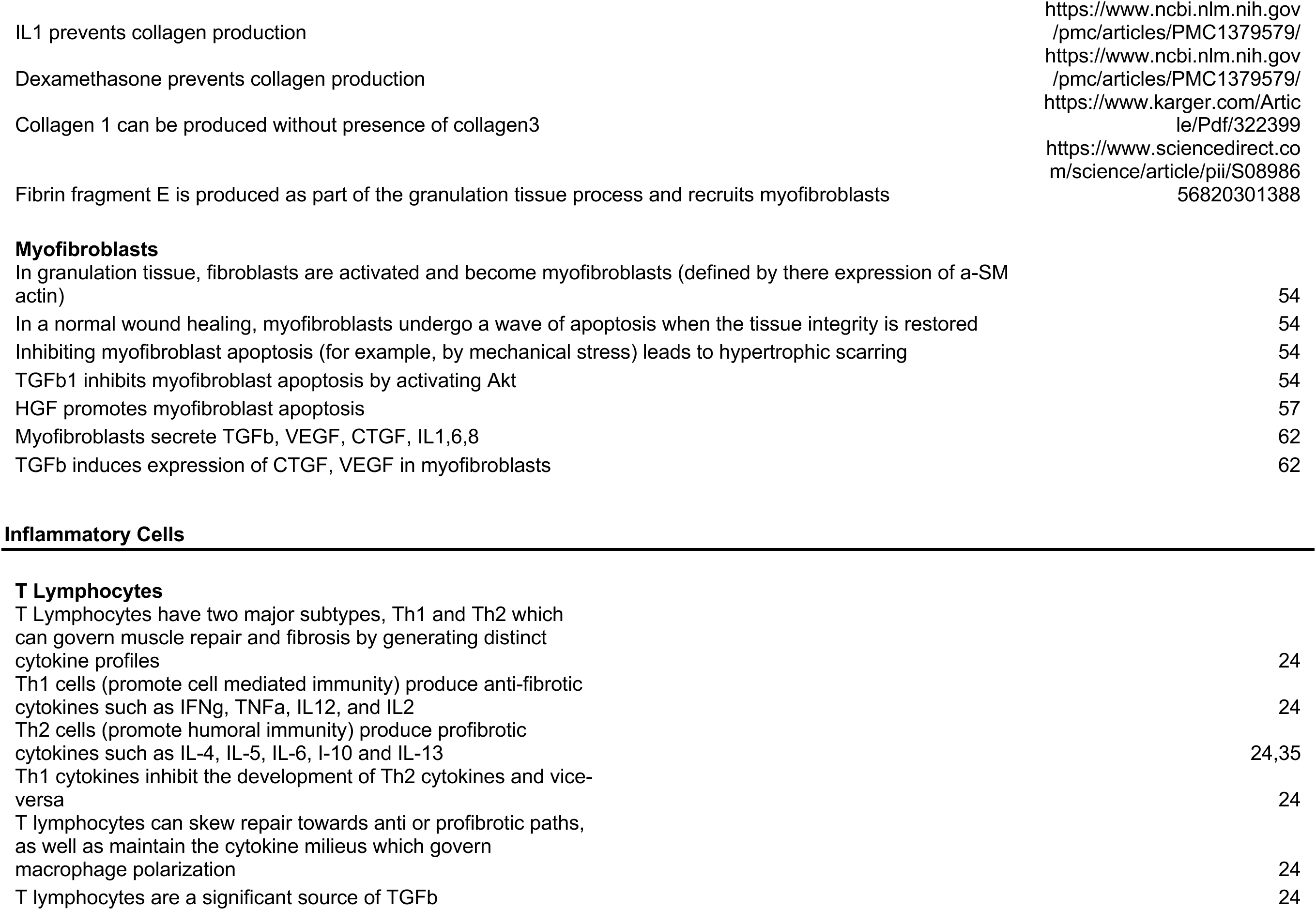

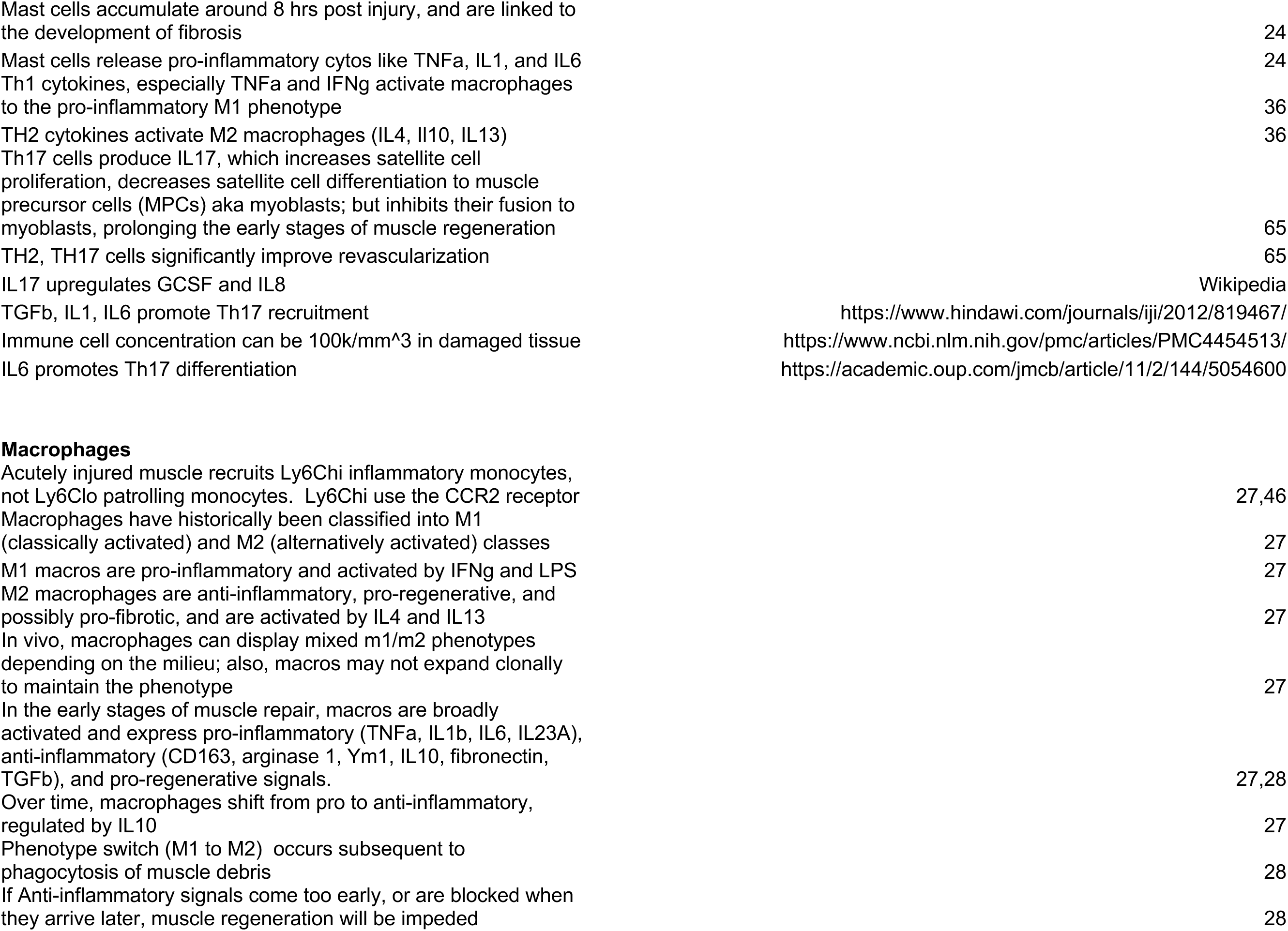

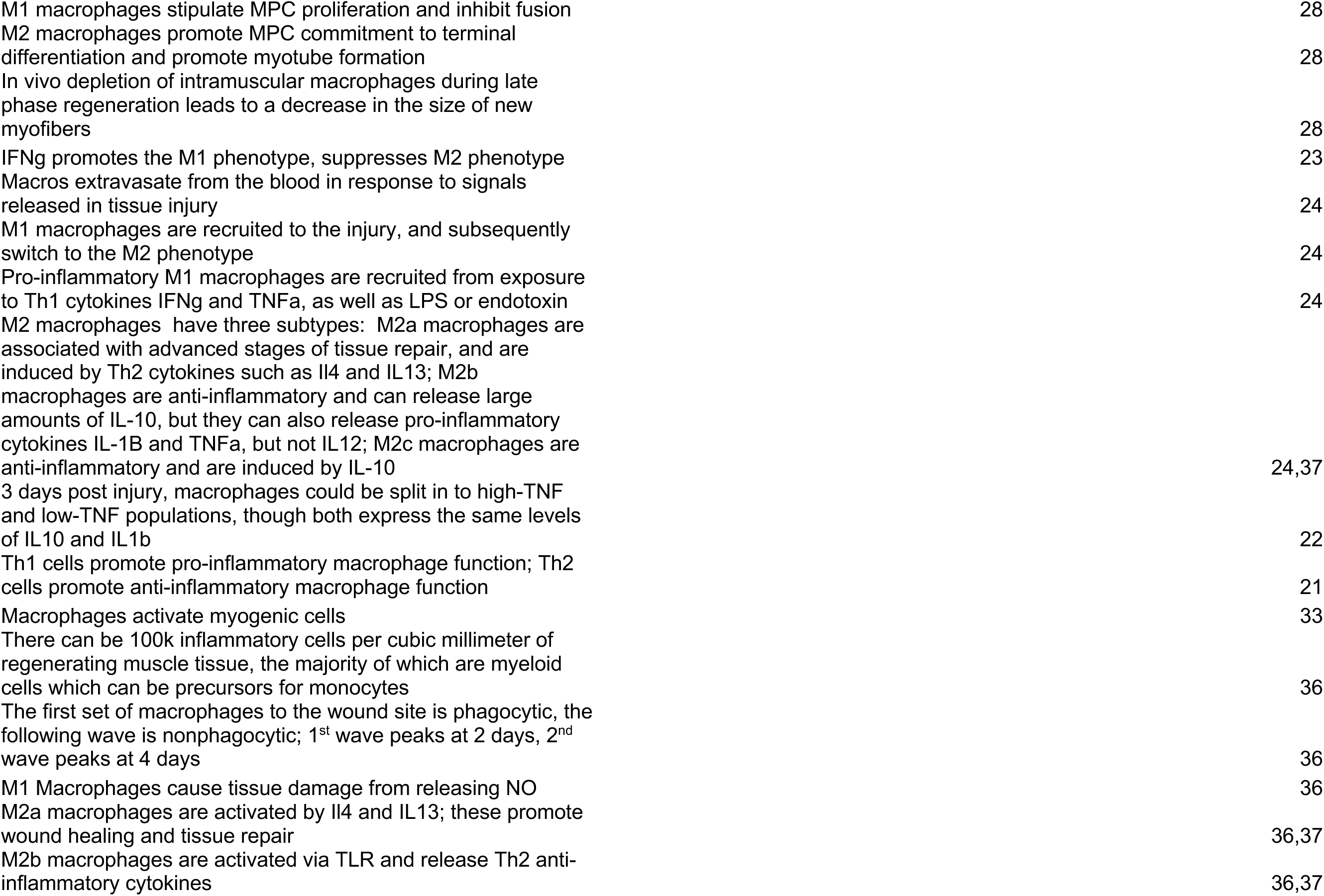

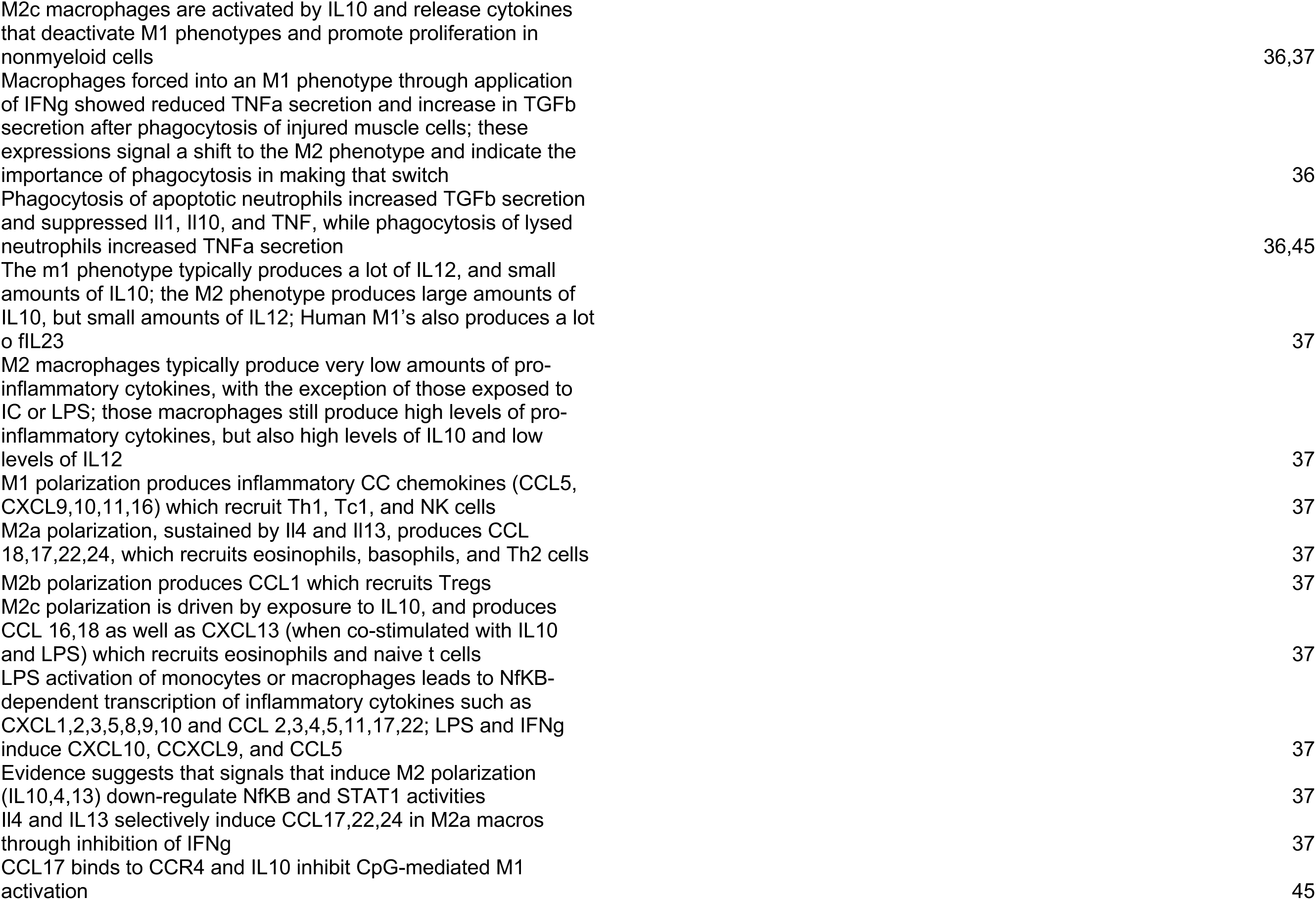

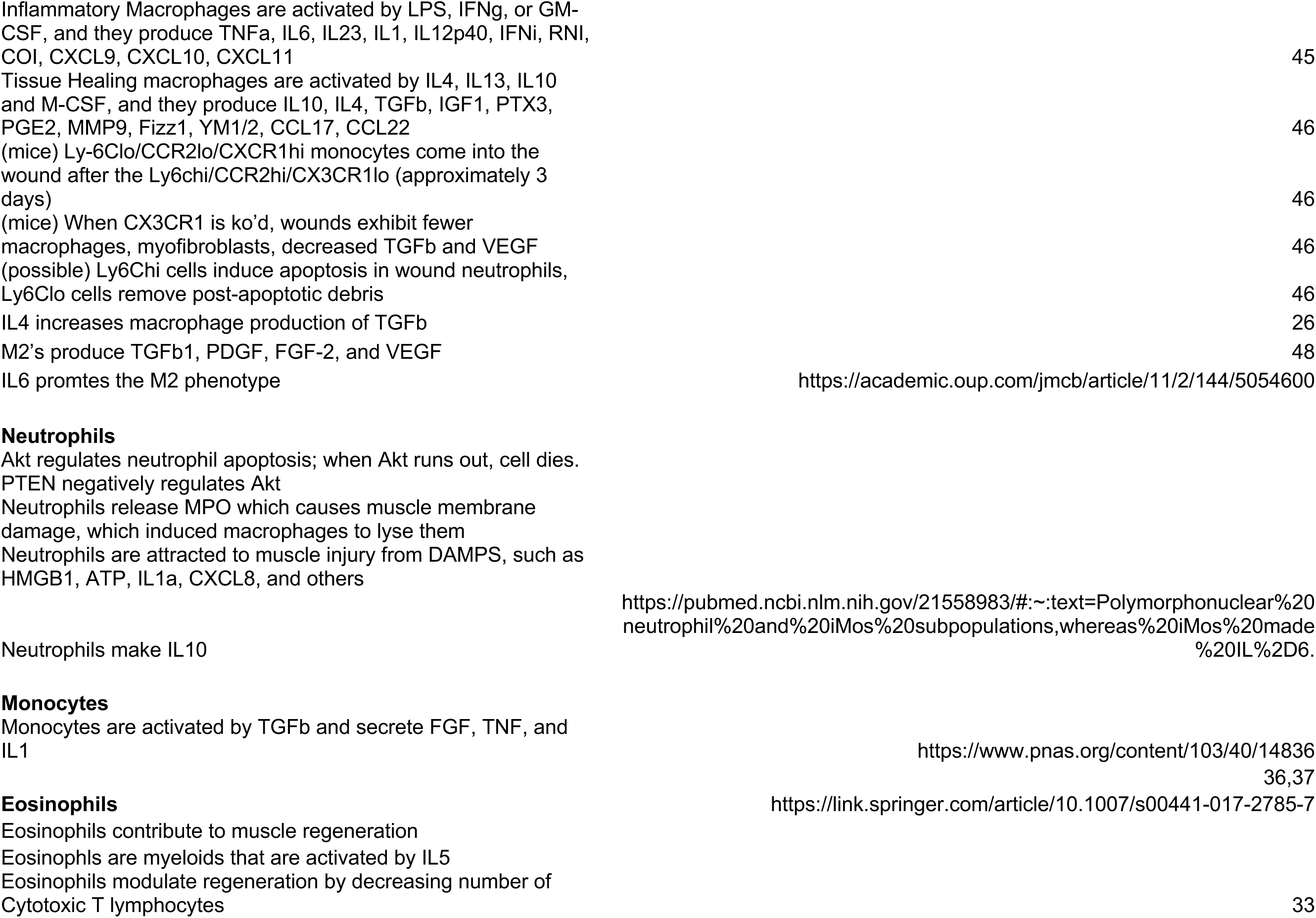

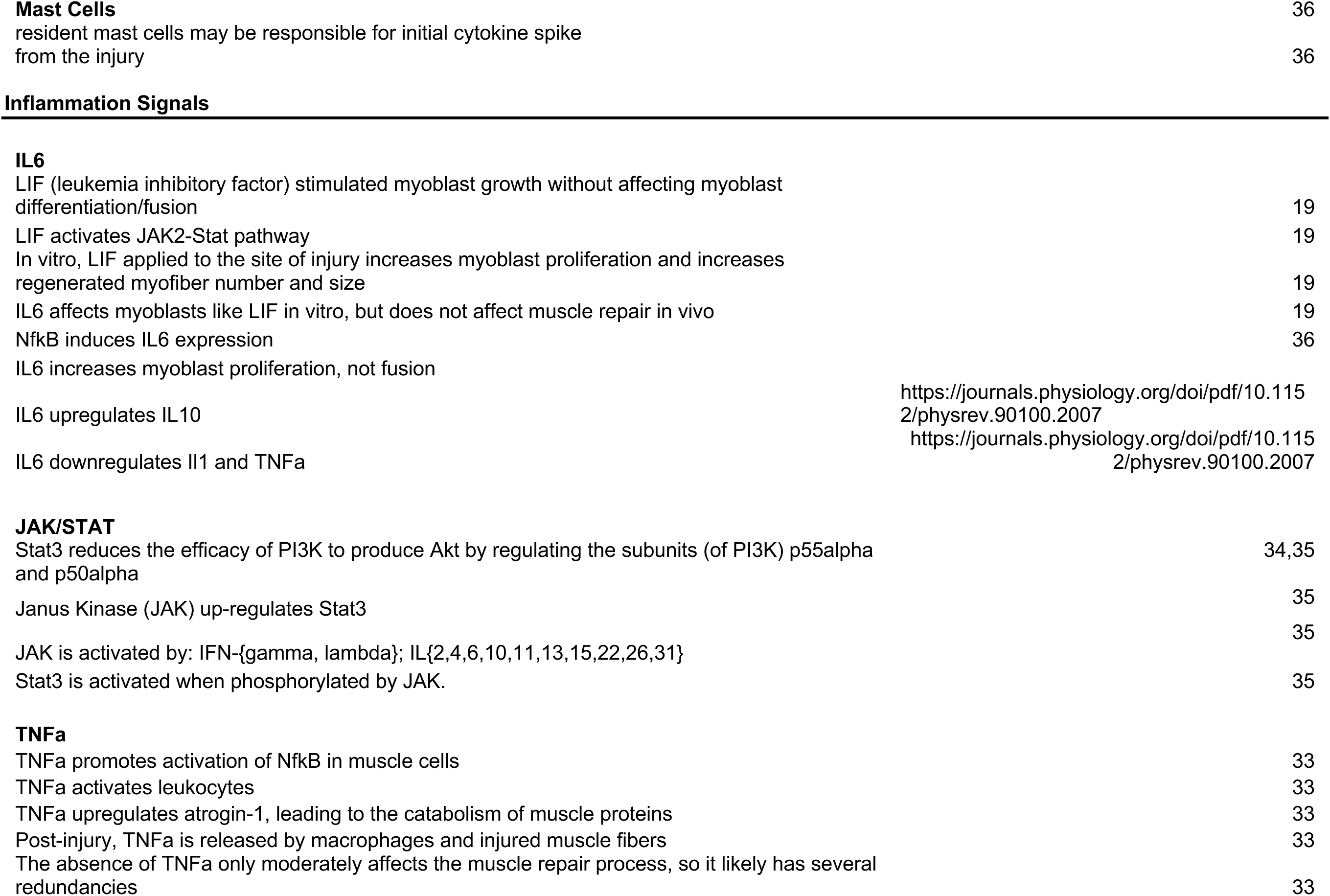

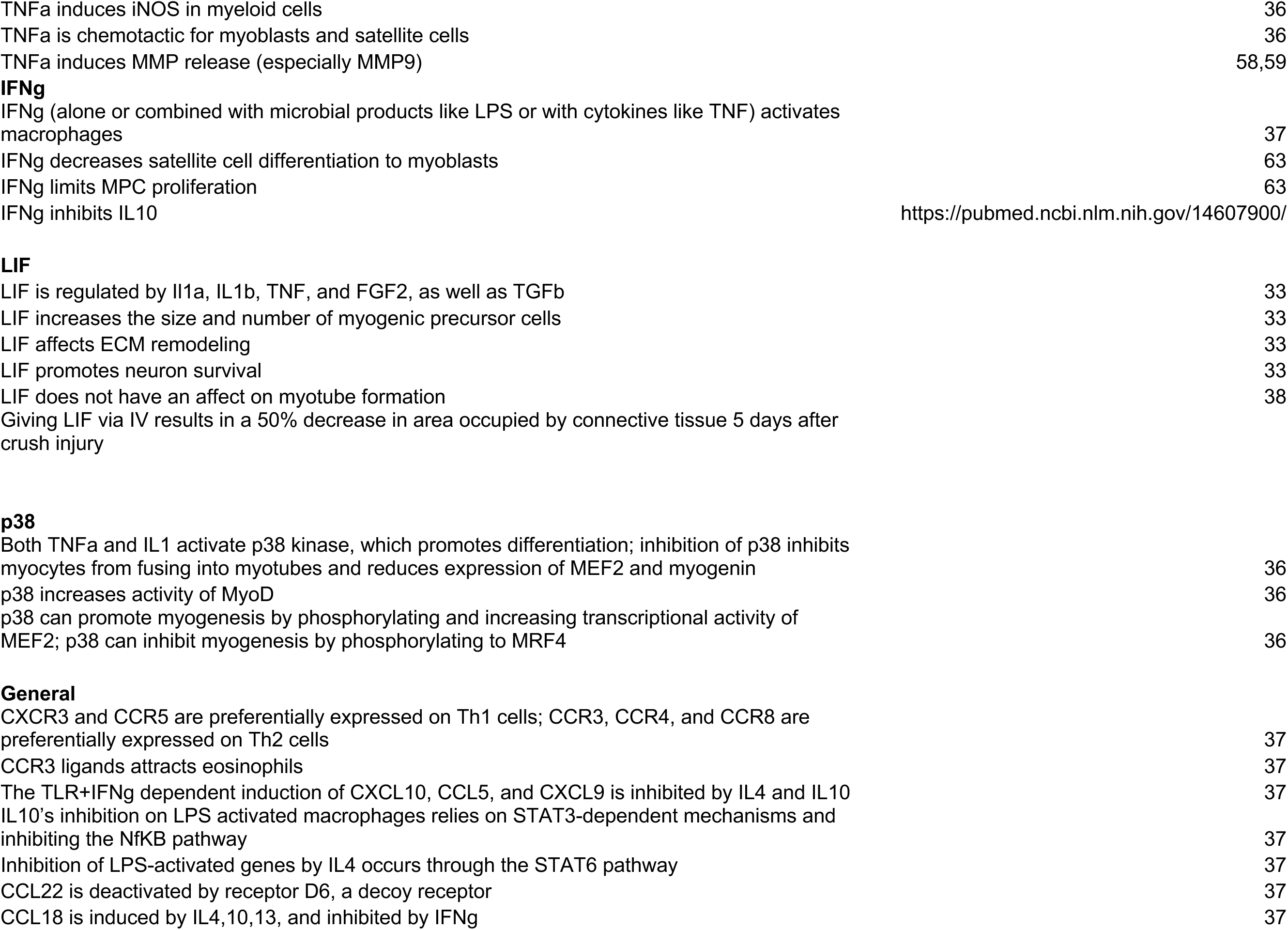

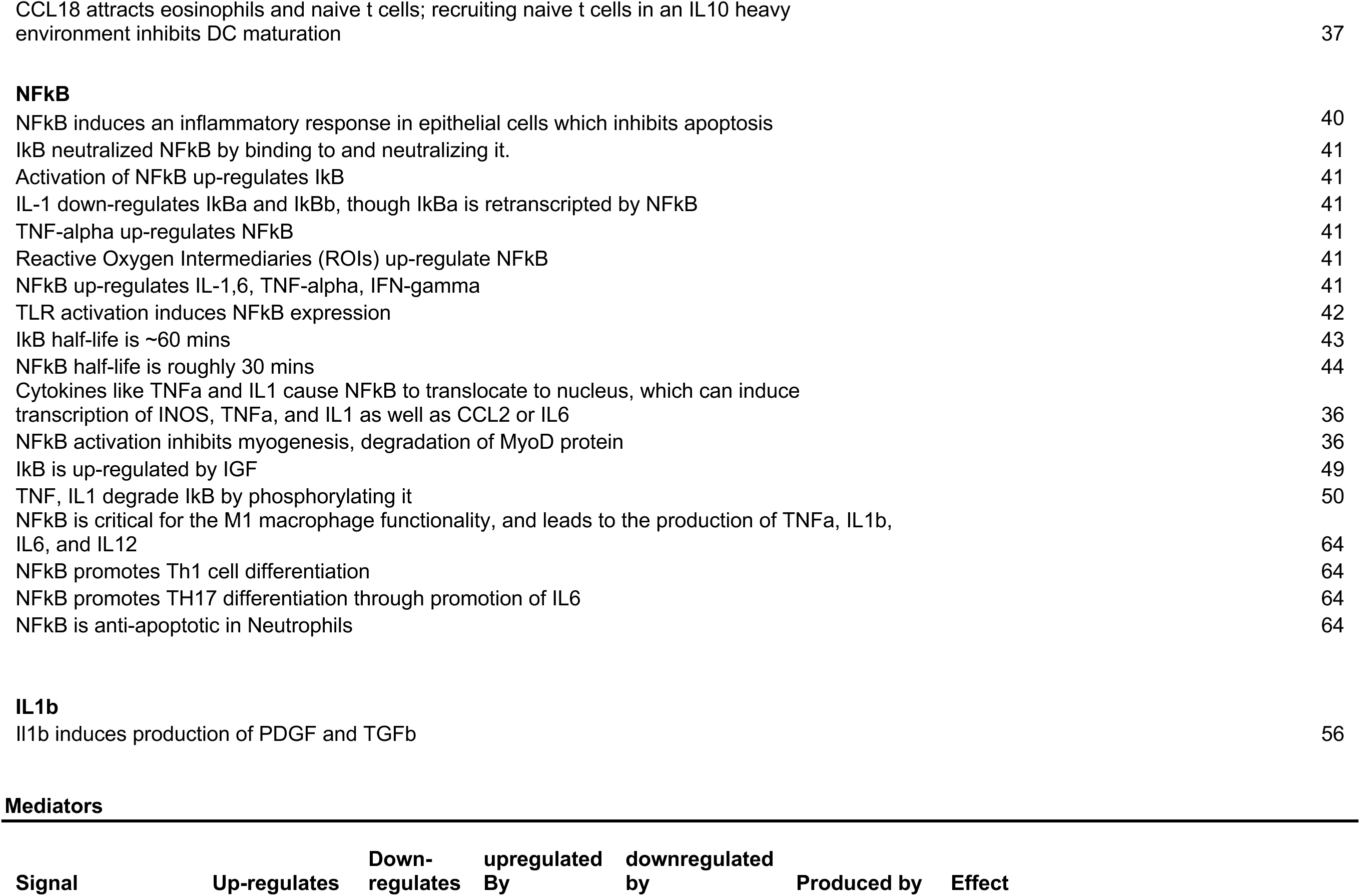

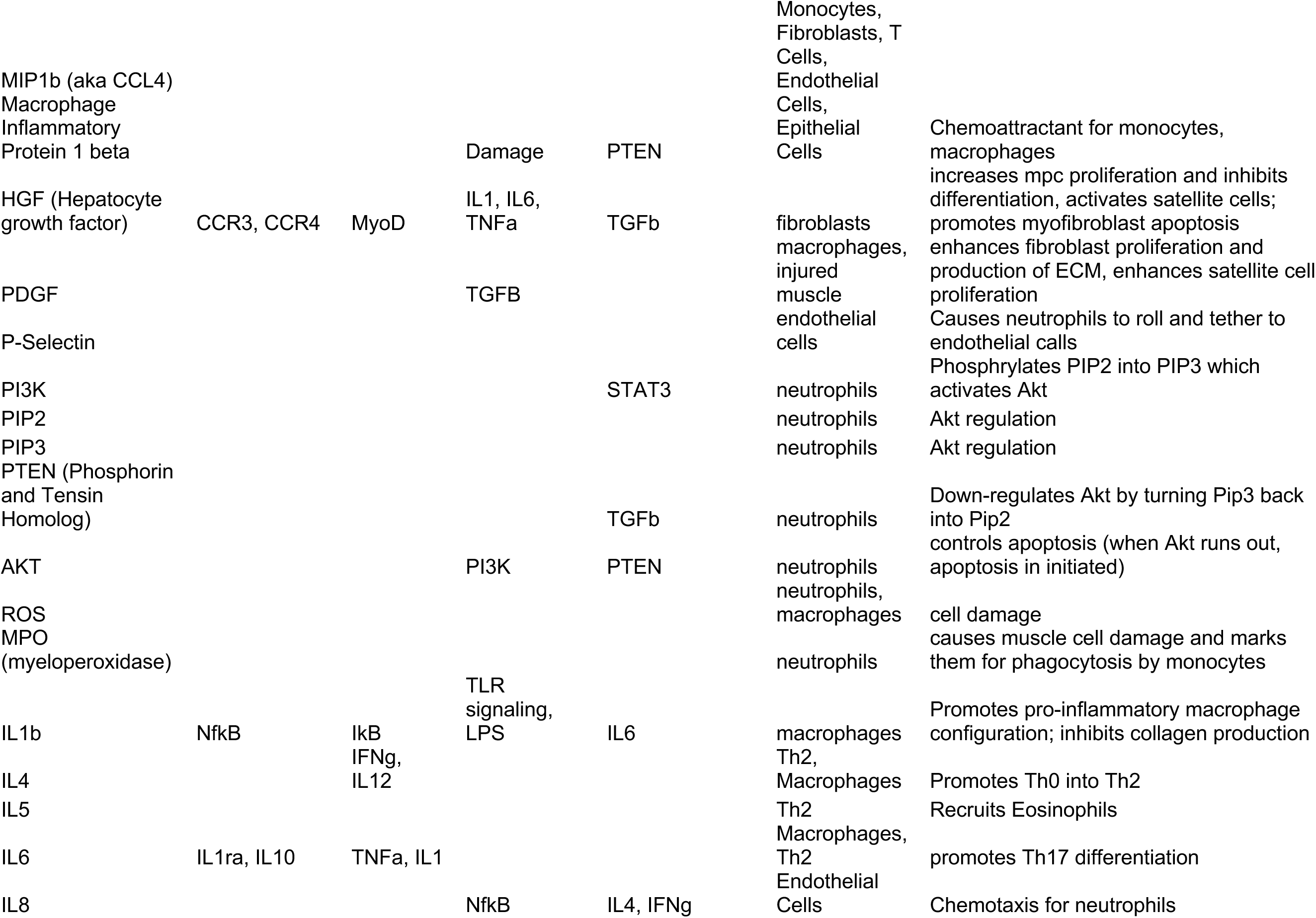

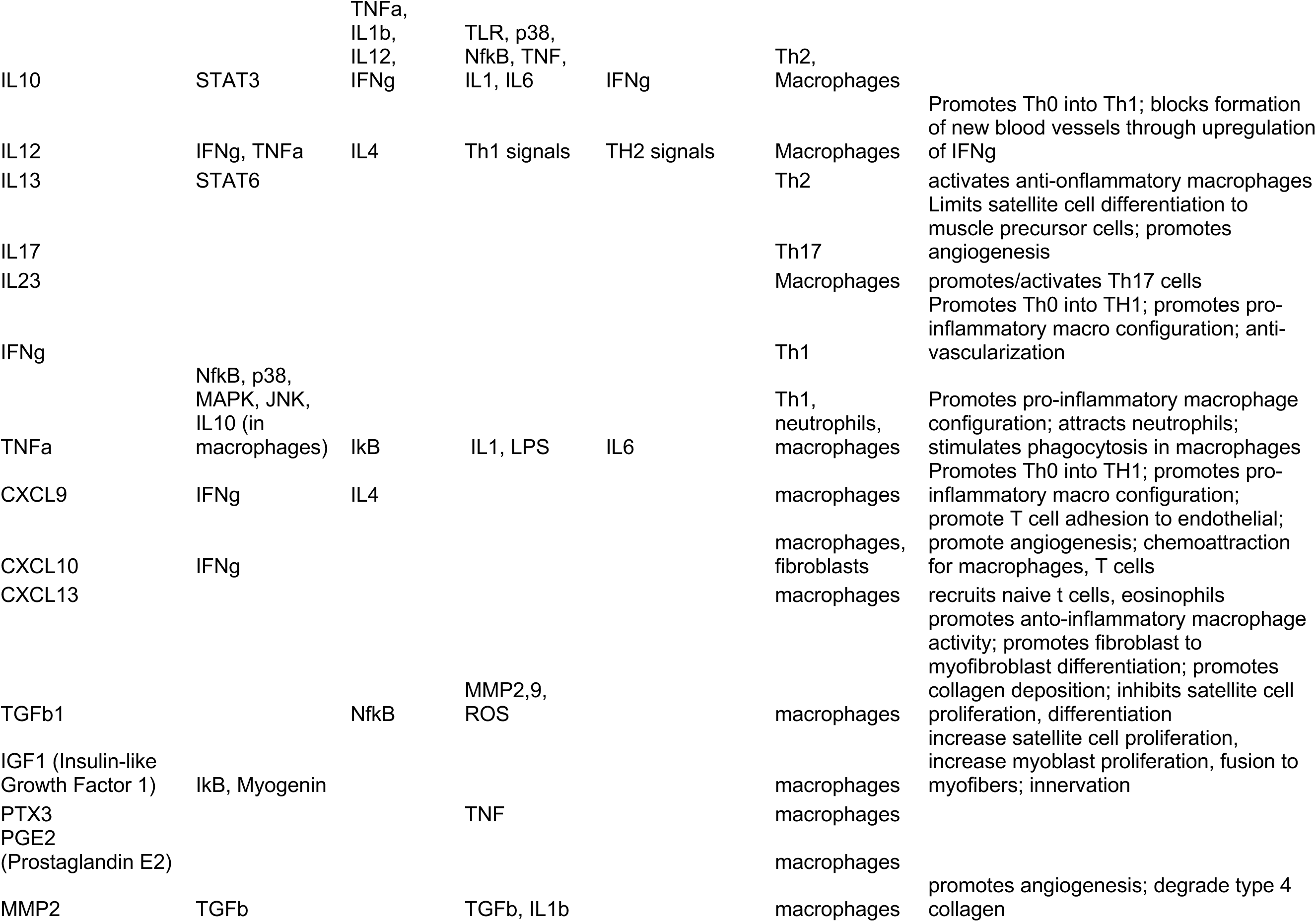

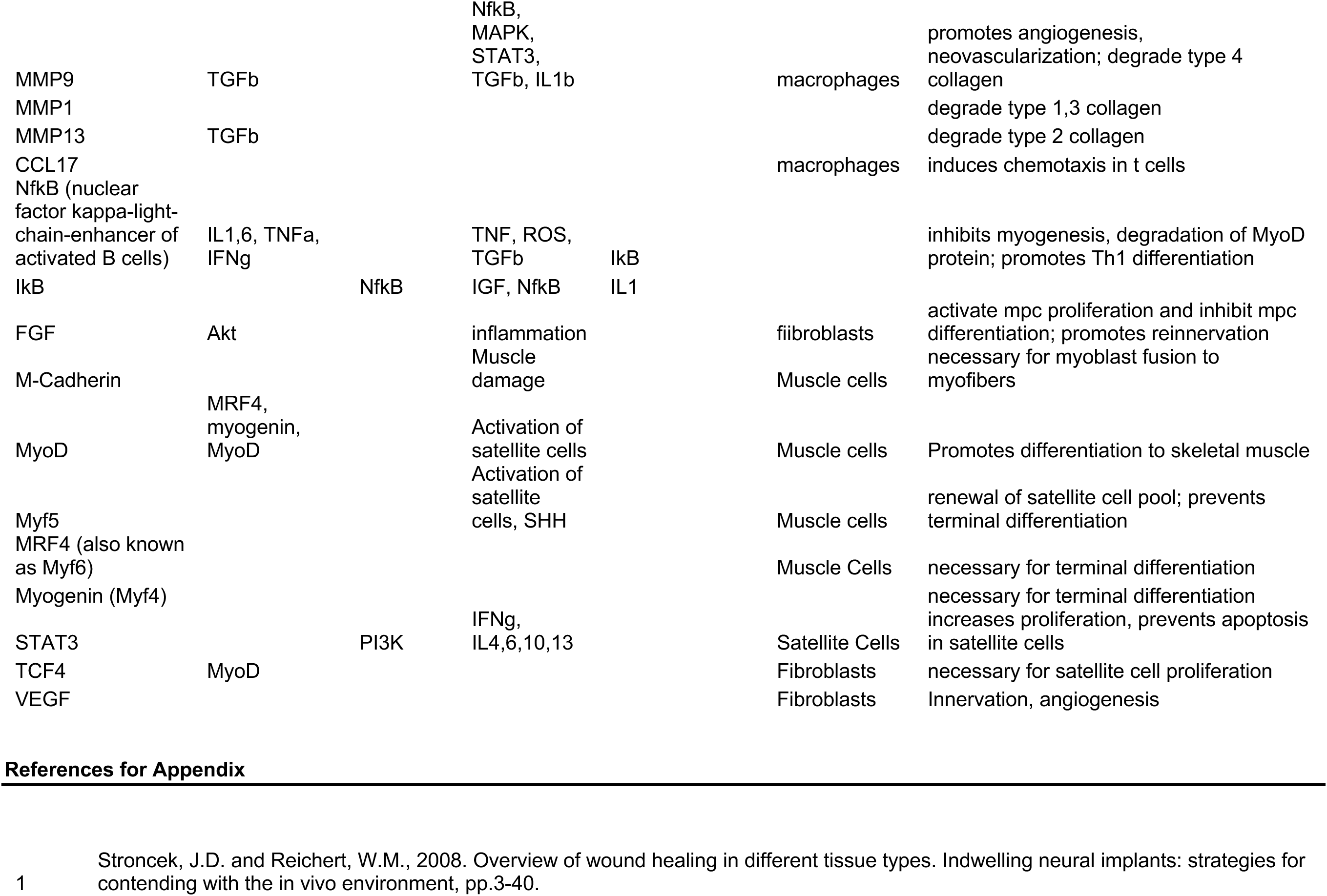

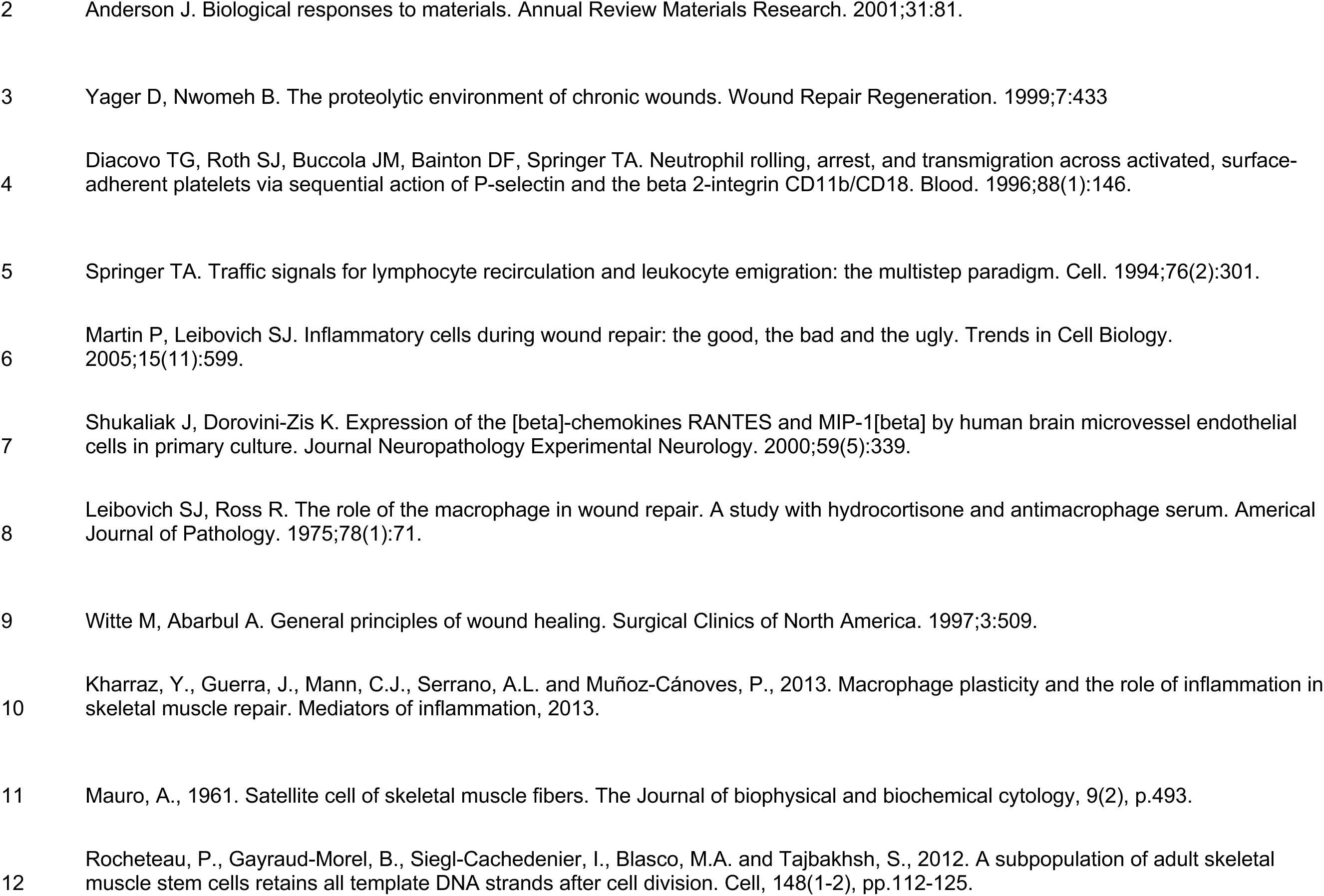

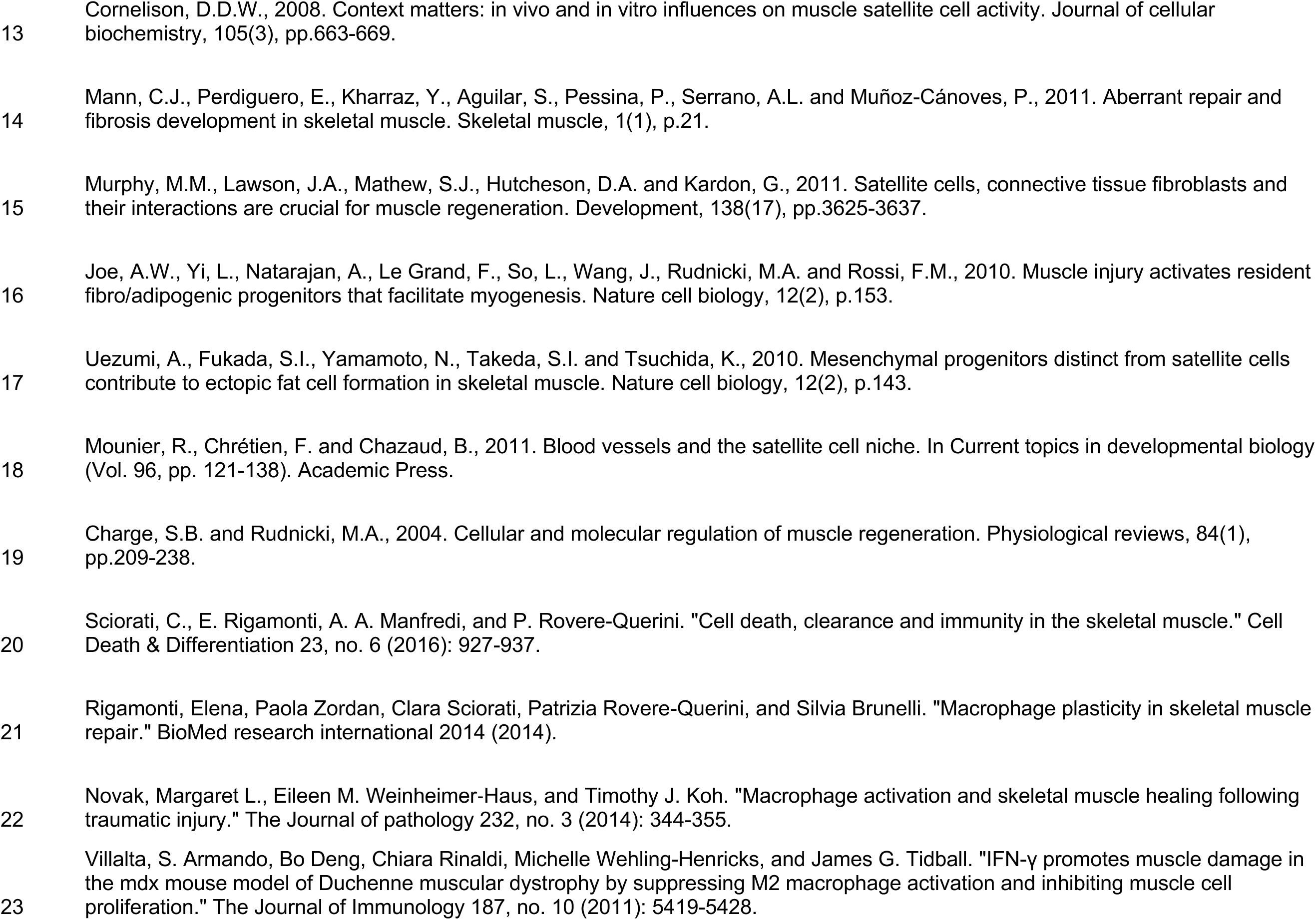

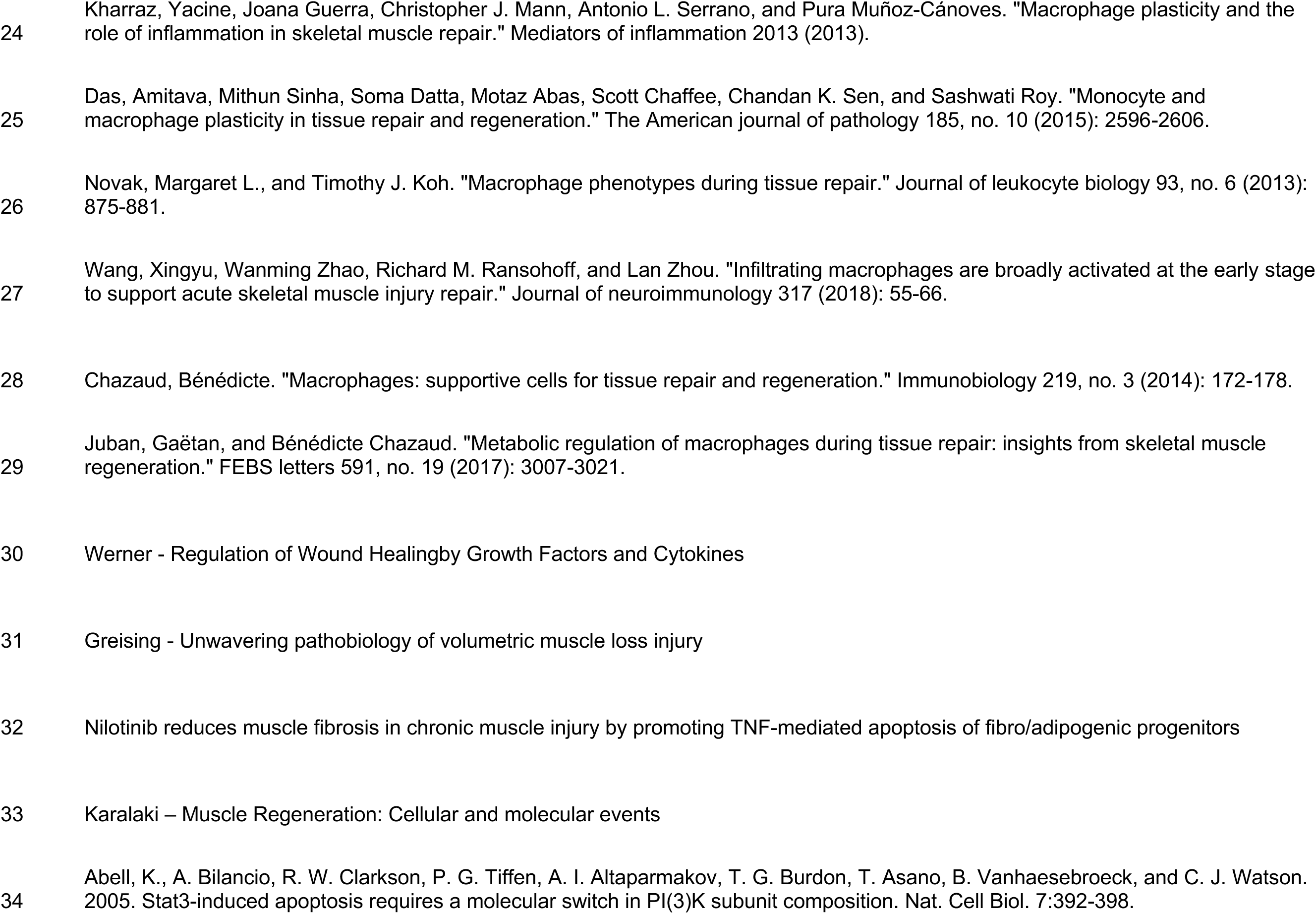

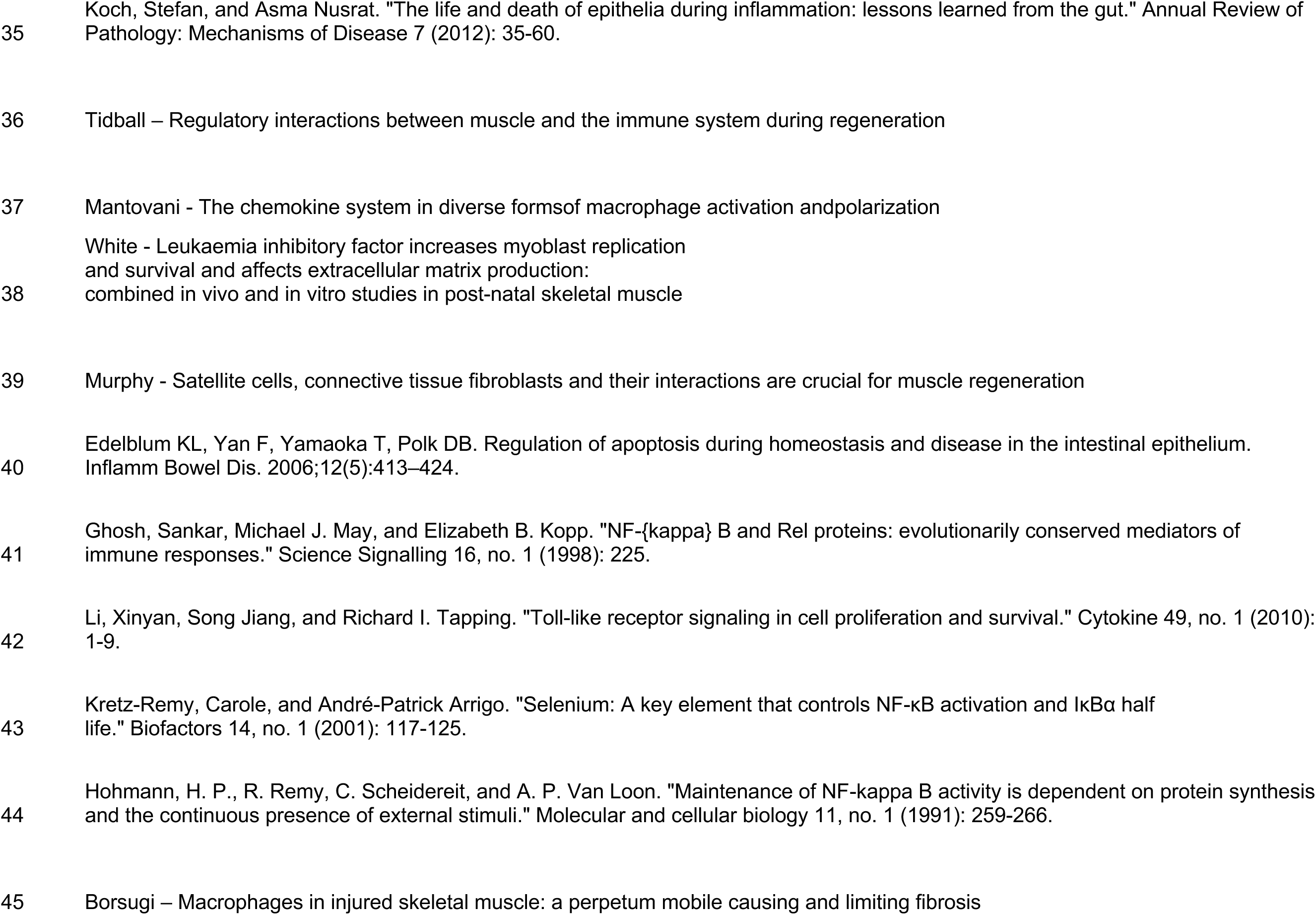

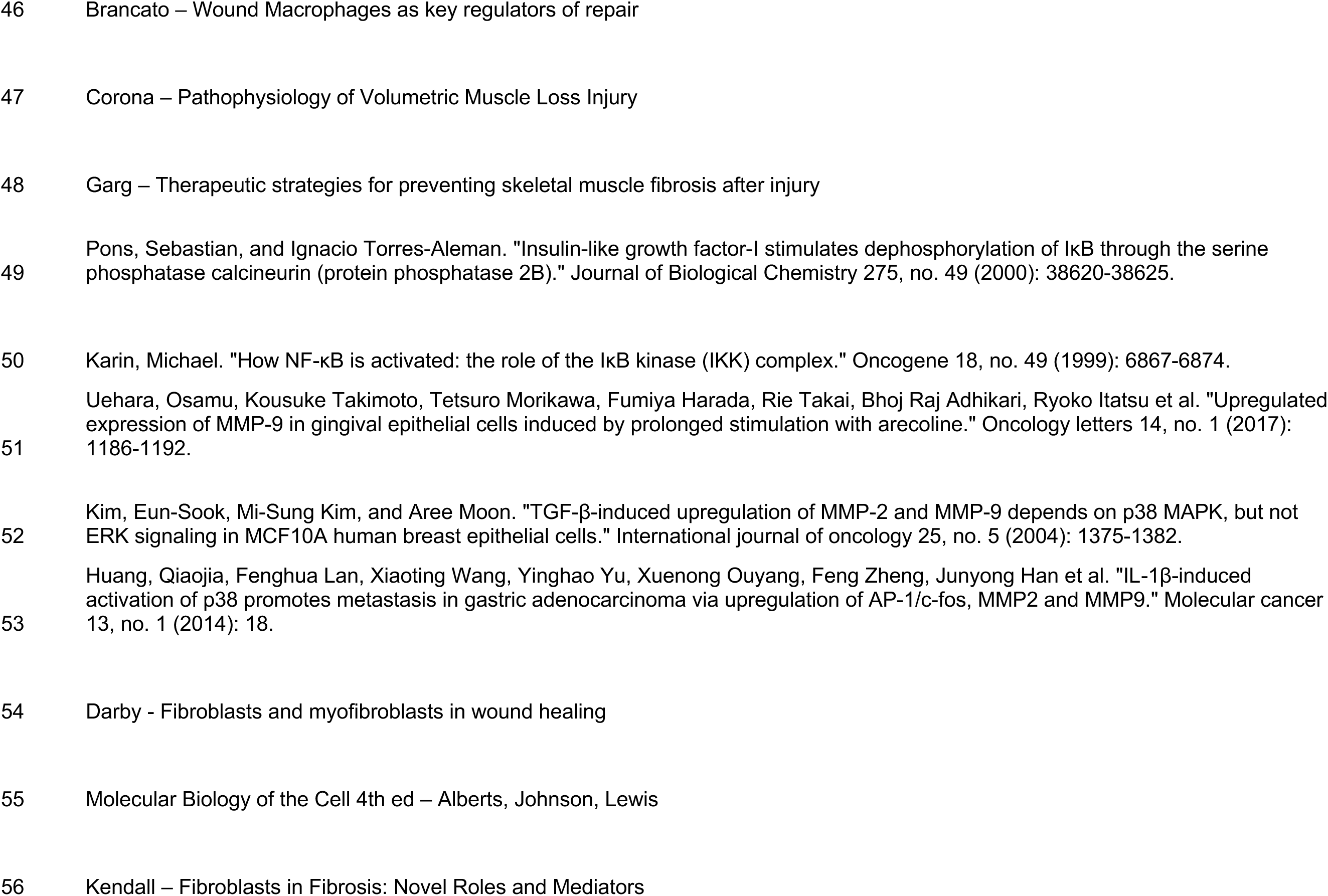

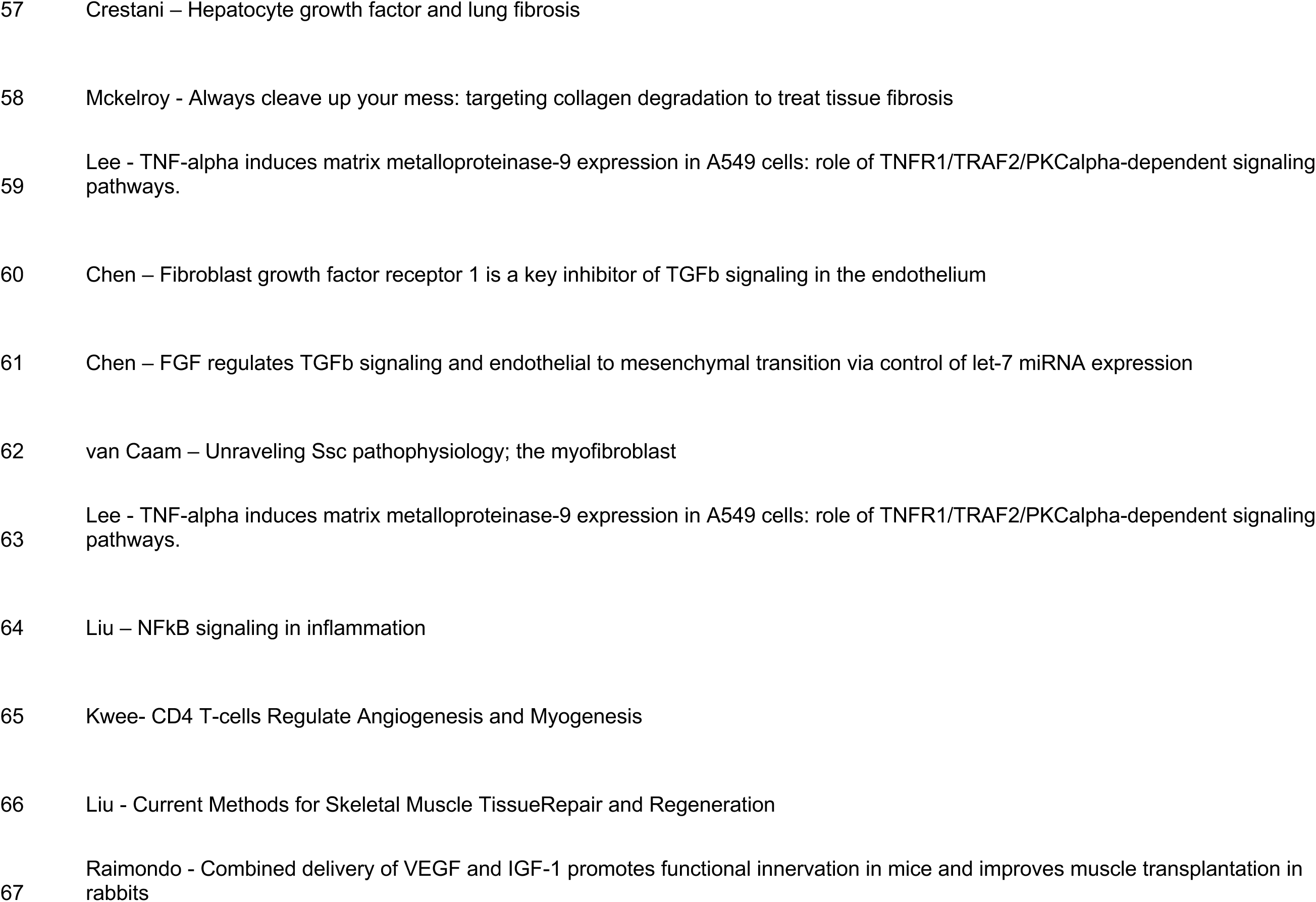

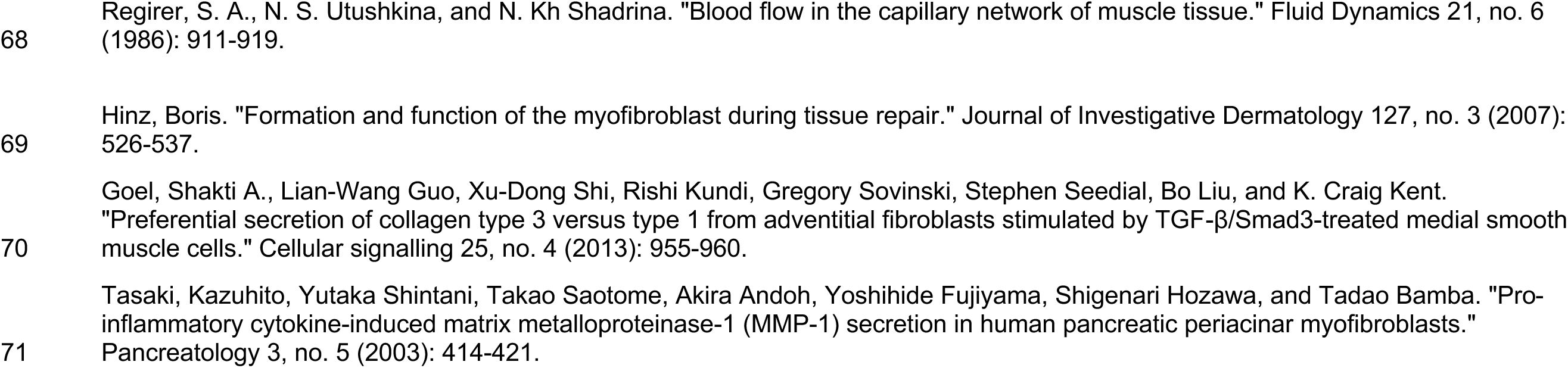

